# Individual reversible plasticity as a genotype-level bet-hedging strategy

**DOI:** 10.1101/2020.05.12.090308

**Authors:** T.R. Haaland, J. Wright, I.I. Ratikainen

## Abstract

Reversible plasticity in phenotypic traits allows organisms to cope with environmental variation within lifetimes, but costs of plasticity may limit just how well the phenotype matches the environmental optimum. An additional adaptive advantage of plasticity might be to reduce fitness variance, or bet-hedging to maximize geometric (rather than simply arithmetic) mean fitness. Here we model the evolution of reaction norm slopes, with increasing costs as the slope or degree of plasticity increases. We find that greater investment in plasticity (i.e. steeper reaction norm slopes) is favoured in scenarios promoting bet-hedging as a response to multiplicative fitness accumulation (i.e. coarser environmental grains and fewer time steps prior to reproduction), because plasticity lowers fitness variance across environmental conditions. In contrast, in scenarios with finer environmental grain and many time steps prior to reproduction, bet-hedging plays less of a role and individual-level optimization favours evolution of shallower reaction norm slopes. We discuss contrasting predictions from this partitioning of the different adaptive causes of plasticity into short-term individual benefits versus long-term genotypic (bet-hedging) benefits under different costs of plasticity scenarios, thereby enhancing our understanding of the evolution of optimum levels of plasticity in examples from thermal physiology to advances in avian lay dates.

**Impact summary:** Phenotypic plasticity is a key mechanism by which organisms cope with environmental change. Plasticity relies on the existence of some reliable environmental cue that allows organisms to infer current or future conditions, and adjust their traits in response to better match the environment. In contrast, when environmental fluctuations are unpredictable, bet-hedging favours lineages that persist by lowering their fitness variance, either among or within individuals. Plasticity and bet-hedging are therefore often considered to be alternative modes of adaptation to environmental change. However, we here make the point that plasticity also has the capacity to change an organism’s variance in fitness across different environmental conditions, and could thus itself be part of – and not an alternative to – a bet-hedging strategy. We show that bet-hedging at the genotype level affects the optimal degree of plasticity that individuals use to track environmental fluctuations, because despite a reduction in expected fitness at the individual level, costly investment in the ability to be plastic also lowers variance in fitness. We also discuss alternative predictions that arise from scenarios with different types of costs of plasticity. Evolutionary bet-hedging and phenotypic plasticity are both topics experiencing a renewed surge of interest as researchers seek to better integrate different adaptations to ongoing rapid environmental change in a range of areas of literature within ecology and evolution, including behavioural ecology, evolutionary physiology and life-history theory. We believe that demonstrating an important novel link between these two mechanisms is of interest to research in many different fields, and opens new avenues for understanding organismal adaptation to environmental change.

## Introduction

Phenotypic plasticity has been recognized as the most important driver of phenotypic change in response to ongoing rapid environmental change and other anthropogenic disturbances (Merilä and Hendry 2014; Fox et al. 2019). Understanding how evolutionary forces shape different types of plasticity, as well as their limitations, can provide important information about how populations may respond to future environmental changes (Charmantier et al. 2008; Scheiner and Holt 2012; Van Buskirk 2012). Reversible phenotypic plasticity allows organisms to adjust their phenotype to better match their current environmental conditions, and can be beneficial when conditions vary within lifetimes (Via et al. 1995; Chevin et al. 2010; Botero et al. 2015; Tufto 2015). However, producing a phenotype that matches the environment perfectly at all times is not always possible, because such phenotypic adjustments come with a wide range of possible costs. Plasticity in traits related to an organism’s morphology or physiology can involve considerable energetic production costs (DeWitt et al. 1998; Van Buskirk and Steiner 2009; Auld et al. 2010). Such costs of plasticity are less striking in labile behavioural traits, but may still involve considerable investment in information acquisition (i.e. sampling) and in maintaining the sensory and cognitive machinery to do so effectively (Stephens 2007). In either case, it is reasonable to assume that increasing degrees of plasticity (the extent to which an organism attempts to more accurately match its environment, or the slope of the reaction norm - see (Dingemanse et al. 2010; Westneat et al. 2015)) come with ever increasing costs, and that an increasingly good match provides diminishing fitness benefit returns. Therefore, the optimal strategy (e.g. the reaction norm slope) in a variable environment may not necessarily involve tracking the environment perfectly by trying to match the phenotype exactly to the current environmental conditions. Instead, under escalating costs and diminishing returns, fitness might often be maximized by exhibiting an optimal, limited degree of plasticity, which avoids the worst phenotypic mismatches with the environment but also avoids the highest costs of plasticity.

In variable environments, optimal strategies do not always maximize arithmetic mean fitness across a range of environmental conditions, but importantly involve a trade-off between mean and variance in fitness. Evolutionary biologists have long recognized that strategies lowering arithmetic mean fitness can be selected for if they also lower genotype variance in fitness, thereby increasing geometric mean fitness (Levins 1962; Lewontin and Cohen 1969; Gillespie 1974). Such strategies are known as bet-hedging (Slatkin 1974; Seger and Brockmann 1987) and have gained a lot of interest in recent years, from theoreticians and empiricists alike (Simons 2011; Starrfelt and Kokko 2012; Crowley et al. 2016; Franch-Gras et al. 2017; Haaland et al. 2019b). Bet-hedging operates at the long-term genotype level, and can thus differ substantially from short-term individual optimization (Yoshimura and Clark 1991; McNamara 1998; Haaland and Botero 2019). Interestingly, this crucial difference between arithmetic versus geometric mean fitness must also apply to any arguments concerning the evolution of optimum levels of plasticity.

Across a wide range of theoretical and empirical studies, phenotypic plasticity is usually only seen as an alternative to bet-hedging (Clauss and Venable 2000; Donaldson-Matasci et al. 2013; Simons 2014; Botero et al. 2015; Furness et al. 2015; Tufto 2015; Grantham et al. 2016; Wang and Rogers 2018; Bonamour et al. 2019; Xue et al. 2019). This is because plasticity evolves in response to reliable cues that predictably convey some information about the current environment that the phenotype needs to match, whilst bet-hedging is an adaptive response to stochastic and unpredictable environmental variation that cannot be tracked (Botero et al. 2015; Tufto 2015). For this reason phenotypic plasticity has rarely been considered as part of any type of bet-hedging strategy (but see Frankenhuis et al. 2013; Scheiner 2013; King and Hadfield 2019; Wright et al. 2019). However, based on the logic of the arguments above, there is nothing to prevent bet-hedging (i.e. additional long-term genotypic benefits from maximizing geometric mean fitness) from acting on the degree of phenotypic plasticity itself, or rather for certain degrees of adaptive plasticity to constitute a form of conservative bet-hedging (*sensu* Haaland et al. 2019b). This is because bet-hedging includes any mechanism that lowers variance in fitness at the price of a lower expected fitness, such as generalist phenotypes (Lynch and Gabriel 1987; Haaland et al. 2020) or traits providing insurance against bad conditions (Poethke et al. 2016; Haaland et al. 2019b). Therefore, the adaptive advantages from bet-hedging could mitigate some of the costs of plasticity whenever improved phenotypic tracking of the environment lowers fitness variance. Here we present a model that shows how the optimal amount of reversible plasticity (e.g. the reaction norm slope) is affected by its bet-hedging effects in fluctuating environments, demonstrating a novel link between these two types of adaptations to both predictable and unpredictable environmental variation.

### Model setup

We consider a simple scenario where some aspect of the environment *e* varies between −1 and 1, with a uniform distribution. An organism can adjust a trait *z* to better match the environment conditions such that the optimal trait value in any given environment is equal to the value of the environmental parameter. We let the ecological payoffs individual *i* gains in any single time step *t* be a Gaussian function,

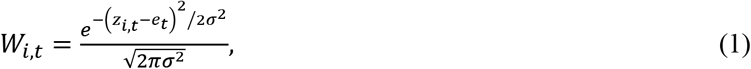

such that payoffs decrease with increasing absolute mismatch between *z* and *e*. *σ* represents the width of the function, and is set to 0.4, making the range of possible *W_t_* values span approximately from 0 (with maximal mismatch) to 1 (with no mismatch, *z_t_-e_t_*=0).

We assume that a cue exists that gives the organism perfect information about the current environmental conditions *e*, and that the organism can exhibit reversible phenotypic plasticity determined by its reaction norm slope *β* in response to the cue, such that the organism’s phenotype in any single time step is *z_t_* = *βe_t_*. An organism perfectly matching the environment in all conditions (i.e. an individual with a reaction norm slope *β* of 1) will then gain equal payoffs in all environments. However, this payoff is limited because the costs of plasticity increase quadratically with reaction norm slope. This represents maintenance costs that do not depend on the number of times the phenotype is adjusted, only on the degree of plasticity itself (see Supporting Information (SI) Appendix B for results with alternative plasticity cost structures). The fitness of an organism *i* after *n* time steps thus becomes:

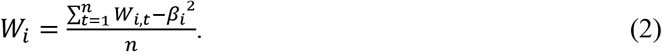

Here we note that individuals with different reaction norm slopes will differ not only in their mean fitness across environments, but also in their variance in fitness (Reed et al. 2010). A non-plastic individual (*β*=0) will incur no costs of plasticity, but will experience high variance in fitness as the environment varies – doing very well under mean environmental conditions (*e*=0), but very badly when environmental conditions deviate a lot from the mean. A perfectly plastic individual (*β*=1) may have low fitness due to its high cost of plasticity, but will experience no variance in fitness across environments, since its phenotypic mismatch is always zero (its phenotype is always ‘equal’ to the environment). Thus, for most scenarios there is likely to be an optimal intermediate reaction norm slope 0<*β\**<1 that best trades off the costs and benefits of plasticity. However, note also that the *β*\* offering the highest arithmetic mean fitness will also provide a non-zero variance in fitness across environments. There may therefore be scope for a bet-hedging strategy involving an additional degree of plasticity, favouring a sub-optimally steeper reaction norm slope *β*>*β*\* (i.e. ‘sub-optimal’ from the point of view of the individual maximizing arithmetic mean fitness across environments), which offers lower arithmetic mean fitness but also a lower variance in fitness. To illustrate this, we simulate 10^7^ realizations of *e*~U(−1,1), and we note that *W* also becomes a random variable, whose properties depend on *β*. Figure 1 shows how different values of *β* provide different arithmetic mean fitness, E[*W*], geometric mean fitness, exp{E[*ln W*]}, and variance in fitness, Var[*W*]. Since Var[*W*] is strictly decreasing with increasing values of *β*, arithmetic mean fitness is maximized by a lower *β* than geometric mean fitness.

**Figure 1:**
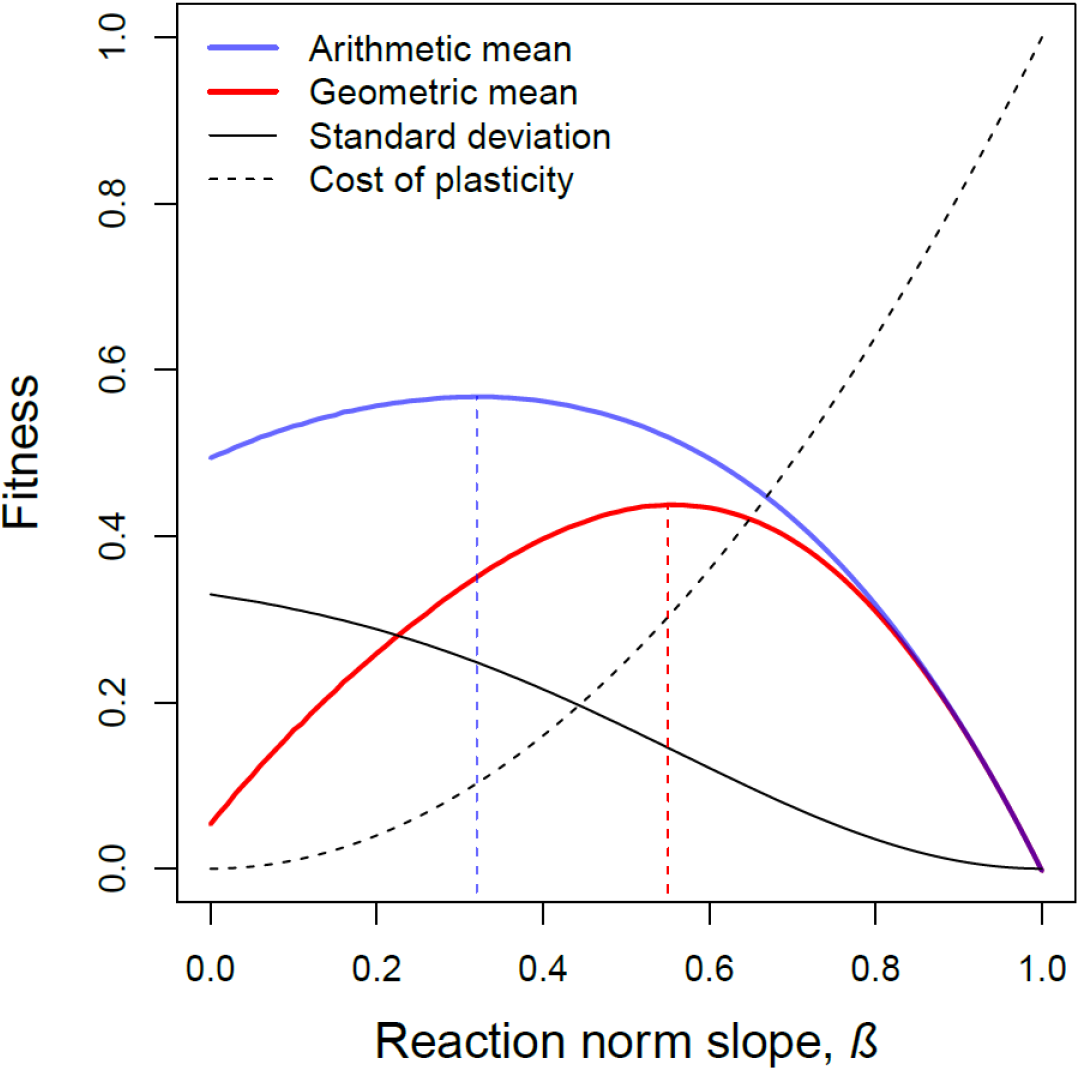
Phenotypic plasticity effects on fitness means and variance. Different values of *β* (reaction norm slope) provide different values for arithmetic (blue) and geometric (red) mean fitness, as well as standard deviation (black) in fitness, across environmental fluctuations. Environments are uniformly distributed between −1 and 1, and fitness in a given environment is given by Eq. 2 (*n* = 1, *σ*^2^ = 0.4). Dashed vertical lines show the maxima of the corresponding functions. Dotted black line: *β*^2^, indicates fitness costs of plasticity. A non-plastic individual (*β*=0) will incur no costs of plasticity, but will experience high variance in fitness as the environment varies – doing very well under mean environmental conditions (*e*=0), but very badly when environmental conditions deviate a lot from the mean. A perfectly plastic individual (*β*=1) may have low fitness due to its high cost of plasticity, but will experience no variance in fitness across environments, since its phenotypic mismatch is always zero (its phenotype is always ‘equal’ to the environment). Since Var[*W*] is strictly decreasing with increasing values of *β*, arithmetic mean fitness is maximized by a lower *β* than geometric mean fitness.

### When should bet-hedging be most important?

By definition, the importance of bet-hedging strategies depends on the extent to which genotype fitness accrues multiplicatively versus additively. An important factor is the correlations in the environmental conditions different individuals experience at a given time (often called the ‘grain’ of the environment, see Starrfelt and Kokko 2012). If all individuals experience the same environment (‘coarse’ environmental grain), fitness accumulates multiplicatively over time, and poor performance in any one environment will be detrimental to lineage survival. Such conditions are typical for many of the best examples of bet-hedging in the real world, such as inter-annual variability in rainfall determining optimal germination strategies for desert annual plants (Cohen 1966; Venable and Brown 1988; Gremer et al. 2012; Gremer and Venable 2014), and timing of dormancy in ephemeral pools for fish (Furness et al. 2015), crustaceans (Gerber and Kokko 2018; Wang and Rogers 2018) and rotifers (Franch-Gras et al. 2017). However, when different individuals of a lineage experience different environmental conditions at a given time (‘finer’ environmental grain), poor performance in any one environment may matter less, since only a subset of individuals are affected, and the lineage may persist through fitness accrued via other individuals experiencing better conditions. In short, among individuals, fitness accumulates additively over space, but multiplicatively over time (Dempster 1955; Frank and Slatkin 1990; Hansen 2018).

Another factor modifying the extent to which a given trait is subject to selection for bet-hedging is how often the trait is expressed within an individual’s lifetime (see Haaland et al. 2019a), and this is especially relevant in the case of reversible plasticity. If the trait is only expressed once per lifetime, fitness accumulation across distinct events only occurs among individuals. For example, in coarse-grained environments, traits relating directly to reproductive success (such as timing of reproduction) in semelparous annual or short-lived organisms should maximize geometric mean fitness across all possible environmental conditions (since total failure in a very early/late year means the whole lineage goes extinct). Such traits should therefore be subject to evolution of adaptive bet-hedging strategies (Haaland et al. 2019a). On the other hand, traits expressed several times under a series of different environmental conditions within a lifetime can accumulate fitness additively across events. For example, the payoffs of a given foraging strategy typically accumulate over a long series of foraging bouts with variable prey availability, and the optimal strategy is therefore assumed to be the one with the highest arithmetic mean energy intake (or energy intake per time) across the sequence of events (Elner and Hughes 1978; Parker and Maynard Smith 1990; Ydenberg et al. 2007). The same can apply to decisions more directly linked to reproductive strategies in iteroparous longer-lived species. If an organism can expect to have many breeding attempts in its lifetime, reproductive success accumulates additively across breeding attempts, and longterm genotype fitness is less affected by reproductive failure in a single year, as long as its strategy works well in most years on average (Haaland and Botero 2019). Therefore, these traits should be seen as maximizing arithmetic mean fitness and involve less scope for bet-hedging.

We therefore simulate the evolution of phenotypic plasticity in scenarios that differ in the grain of the environment (correlations among environments experienced by different individuals at a given time step) and in the number of times per lifetime the potentially plastic trait contributes to accumulating fitness prior to selection. As explained above, we expect bet-hedging to favour apparently sub-optimally steeper reaction norms in scenarios with coarser environmental grains and fewer distinct events across which fitness is accumulated.

### Evolutionary simulations

Our individual-based simulations follow the evolution of a gene *β* affecting the degree of phenotypic plasticity in a fitness-determining trait. At each time step *t* within a season, an overall environmental condition *E_t_* is determined by a random draw from a continuous uniform distribution between 0 and 1. The microenvironmental conditions experienced by each individual *e_i,t_* can then deviate randomly from *E_t_* depending on the environmental grain *g*. Each *e_i,t_*, value is drawn from a uniform distribution with a lower bound of *e_min_* = *gE_t_* and an upper bound of *e_max_* = *E_t_*+(1-*g*)(1-*E_t_*), such that when *g* = 1 *e*_min_ = *e*_max_ and all individuals experience the same conditions (*E_t_*). When *g* = 0, *e*_min_ = 0 and *e*_max_ = 1, and the overall environment does not affect the distribution of individually-specific microenvironments at all. Finally, we transform all microenvironments with 2*ei*,*t*-1, to ensure that the mean environmental conditions are 0.

Each individual’s phenotype in a given time step, *z_i,t_* is determined by its personally experienced microenvironment and its gene value for phenotypic plasticity (i.e. reaction norm slope), and so *z_i,t_* = *e_i,t_ β_i_*. The fitness gained in that time step is then a Gaussian function of the absolute difference or mismatch between *z_i,t_* and *e_i,t_*. After *n* time steps, individual fitness values *W_i_* are given by equation 2, which is their arithmetic mean fitness across time steps, including the cost of plasticity. Then all individuals reproduce, and their number of offspring produced is equal to each individual’s *W_i_* (on a scale between 0 and 1), multiplied by a constant *r* representing maximum reproductive success. Population size varies with the number of offspring produced, but cannot exceed the carrying capacity *K*. Recruited offspring inherit their parent’s gene value for *β*, or (with probability *m*=0.005) a mutated value drawn from a Gaussian distribution around the parent’s *β* with standard deviation *m_δ_*. We investigate scenarios with both discrete generations, where all adults die at the end of a season and the next generation is made up entirely of offspring produced in that season, and with overlapping generations, where a proportion *α* of the adult population (randomly selected) dies and the population the next season is made up of a combination of surviving adults and newly recruited offspring.

Populations are initiated at carrying capacity, with gene values for *β* uniformly distributed between 0 and 1. We allow the simulations to run for 2000 years, and replicate each parameter combination 100 times to assess consistency of the evolutionary outcomes. All simulations were run in R version 3.6.0 (R Core Team 2019) and code is provided in SI Appendix C.

## Results

Figure 2 shows the main result for simulations with overlapping generations, and very similar results were obtained from simulations with discrete generations (Fig. S1). Evolutionary trajectories quickly stabilized and outcomes were highly consistent across replicate simulations, as shown by the narrow error bars in Fig. 2. As expected, steeper reaction norm slopes are favoured in scenarios expected to promote bet-hedging in response to multiplicative fitness accumulation (coarser environmental grains and fewer time steps prior to reproduction, i.e. high *g* and low *n*), and this is because plasticity lowers variance in fitness across environmental conditions. However, it should be noted that even for scenarios with high *n*, reaction norm slopes continue to decline with lower values of *g*. This is not because of changes in the nature of any fitness accumulation (*n* = 10 already ensures an important role for arithmetic mean fitness within lifetimes), but rather because the amount of temporal environmental variation itself decreases in environments with a finer spatial grain (Fig. S2). In our specific formulation, mean *e_i,t_* tends towards 0 as *g* decreases, even when *E_t_* is near −1 or 1 (since microenvironments *e_i,t_* are bounded by −1 and 1), but such a decrease in temporal variation as spatial variation increases is a common feature of natural environmental variation.

**Figure 2:**
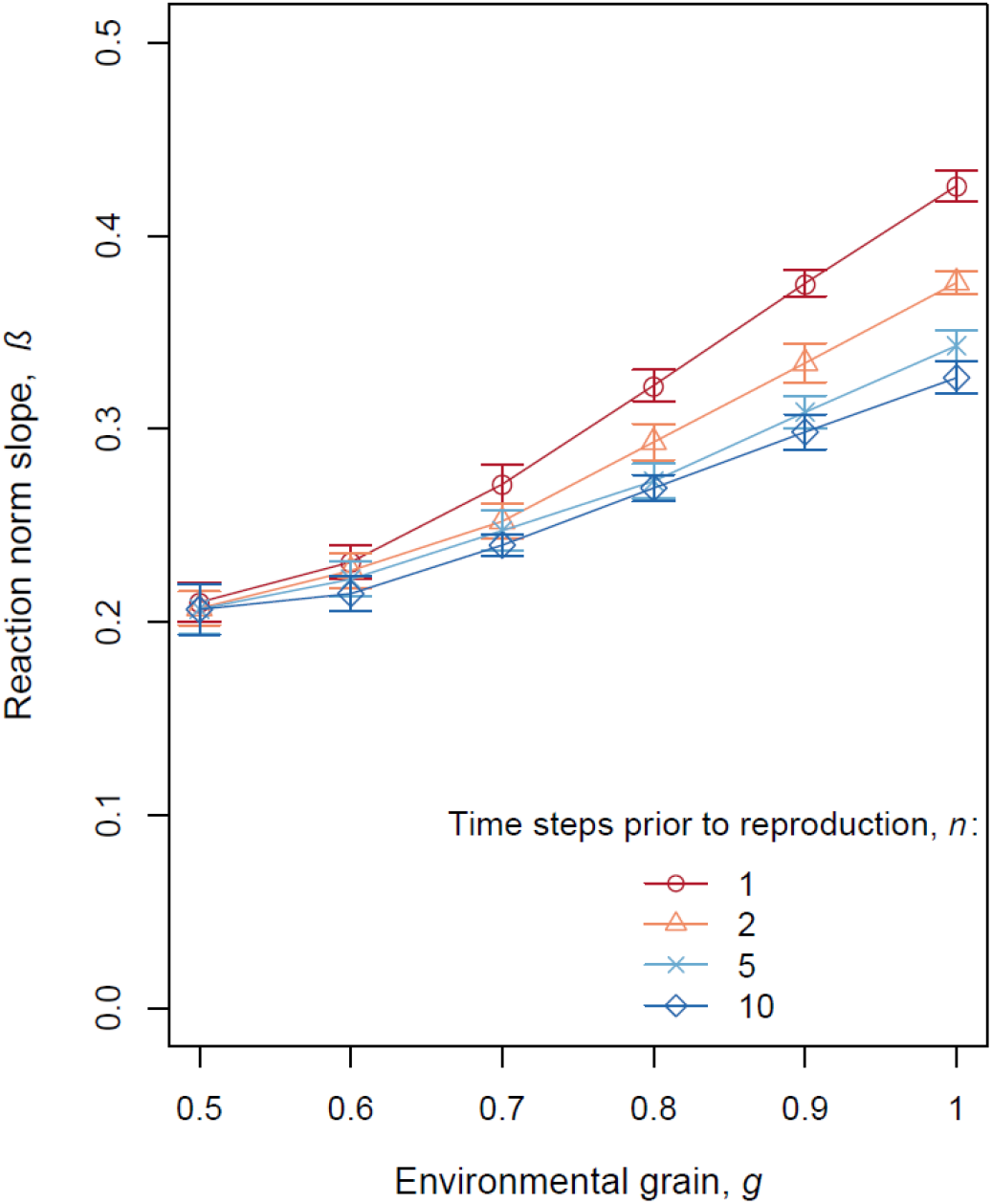
Mean evolved values of phenotypic plasticity *β* (reaction norm slope) from simulated scenarios with overlapping generations (*α* = 0.5) for different values of environmental grain, *g* (x-axis), and number of time steps prior to reproduction, *n* (line colour, point type). Error bars show standard deviations in population means across 100 replicate runs. Baseline parameter values as shown in Table S1.

We observe that for *g* = 1, simulation outcomes correspond well with our predictions from figure 1. Reaction norm slopes in simulations with *n* = 1 evolved to the value maximizing geometric mean fitness (evolved 0.425, predicted 0.43), and in simulations with *n* = 10 they evolved towards the value maximizing arithmetic mean fitness (evolved 0.326, predicted 0.32). As *g* decreases (finer environmental grain), the difference in evolutionary outcomes between high and low *n* scenarios also decreases (figure 2).

In a second set of models (See SI Appendix B), we model an alternative scenario where the cost of plasticity is relative to the amount of phenotypic change from one time step to the next. This ‘production’ cost scenario can lead to fitness variance being minimized for shallower than expected reaction norm slopes (Figs. S3A, S4A, S5-6), because of the increasing variance in costs caused by steeper reaction norms changing the phenotype by a greater degree per time step. The slope favoured by geometric mean under this scenario can thus be shallower than the slope favoured by arithmetic mean. However, when the costs of plasticity include both a general cost relative to the reaction norm slope (as above) *and* such ‘production’ costs (based upon the absolute amount of phenotypic change required each time), bet-hedging selection maximizing geometric mean fitness tends to favour steeper reaction norm slopes, as in our main results (see Figs. S3B and C, and S4B and C).

## Discussion

When fitness accumulates multiplicatively, as occurs in coarse-grained environments for traits that are expressed only once or a few times per lifetime (Starrfelt and Kokko 2012; Haaland et al. 2019a), adaptive bet-hedging at the genotype level can cause individuals to invest more in phenotypic plasticity than they would do otherwise. This is because although this additional plasticity comes at a cost to their expected reproductive output, it also lowers their variance in fitness, and is therefore selected for at the genotype level. This increased investment in phenotypic plasticity represents a type of what has been termed conservative bet-hedging (see Haaland et al. 2019a), since it lowers genotype fitness variance by lowering fitness variance at the individual level. Essentially, the genotype benefits from individuals ‘playing it safe’ and being able to cope better with a wider range of different environments than simply benefits them as individuals. Note that the same tension between individual-versus genotype-level fitness interests is seen in the alternative and more widely recognized diversifying bet-hedging strategy (Cohen 1966; Philippi and Seger 1989), which ‘spreads the risk’ among individuals, such that variance in fitness remains high at the individual level, but is lowered at the genotype level. Under diversifying bet-hedging some individuals are essentially ‘sacrificed’ for the good of the genotype. Hence, under a conservative bet-hedging strategy, such as the type evolving here via additional plasticity, individuals do not necessarily maximize their expected (arithmetic mean) fitness over their lifetimes, if this is selected against at the long-term genotype level in order to minimize any variance in their fitness (Simons and Johnston 2003; Starrfelt and Kokko 2012).

The main result here that traits contributing to fitness fewer times within a lifetime show more plasticity (due to conservative bet-hedging) may explain certain among-species patterns in plastic adjustment in breeding phenology in response to recent climate change (Radchuk et al. 2019). Across generations, traits directly related to reproductive decisions contribute purely multiplicatively to fitness in semelparous species (that reproduce only once), but also additively in iteroparous species, reducing the importance of bet-hedging as the expected number of breeding opportunities per lifetime increases (Haaland et al. 2019a). For example, fluctuations in spring phenology lead to different optimal breeding dates among years, and a lot of recent research has attempted to disentangle the relative roles of plasticity and genetic evolution in reducing the phenological mismatch among optimal and actual timing of breeding, in order to better assess how earlier spring onset due to anthropogenic climate change may threaten population viability (Visser et al. 2004; Charmantier et al. 2008; Tarka et al. 2015; Bonamour et al. 2019). Using data from a recent meta-analysis across a range of bird species (Radchuk et al. 2019) we found, rather counterintuitively, that species with shorter lifespans and fewer expected breeding attempts per lifetime are significantly more likely to show an adaptive plastic response to phenology than species with longer lifespans (see Box 1). In line with our predictions, the longer-lived species in such comparisons may be less likely to evolve to pay the cost of plastically adjusting the timing of their breeding, because experiencing among-year variation in reproductive success is less harmful when there is a higher probability that one can simply try again next year.

### Box 1: A role for bet-hedging in plastic phenological responses to spring temperature

Radchuk et al. (2019) identified estimates of phenological responses to temperature variation across 14 species (their Fig. 3, responses in black). Here we extend their analyses and test whether the number of breeding seasons a sexually mature adult can expect to experience in its lifetime (i.e. the number of times a trait contributes additively to fitness) affects the magnitude of responses in spring phenology to temperature. We calculated this assuming a constant adult survival rate, except in the case of roe deer (*Capreolus capreolus*), where age-specific survival estimates were available (Plard et al. 2014). Fig. B1 shows the result of a log-linear mixed-effects model, which reveals a significant negative effect of expected number of breeding seasons per lifetime (x-axis) on the reaction norm slopes of phenological responses to temperature (y-axis). The slope of the regression line is −0.178 ± 0.095. Population is included as a random effect in the model, since most studies report multiple estimates (predictor-response variable combinations) for the same population. Estimates are weighted by the inverse of the standard error of the mean (point size). Including species or publication ID as random effects did not affect the magnitude of the response (see Table S2, SI Appendix A).

Note that the final species in fig. 3 in Radchuk et al. (2019), the common guillemot *Uria aalge*, has a long lifespan yet shows a strong phenological response. The responses of this colonial seabird are in the opposite direction of the others included in the meta-analysis, and represent a substantial outlier for our current model with absolute reaction norm slopes. We have chosen to omit this single data point because, as the authors of the study in question point out (Reed et al. 2009), the responses to environmental factors are complex and non-linear, and additional strong social pressures are also at play favouring synchronous reproduction in breeding colonies. Including this data point weakens the significance, but the trends remain strong (Table S2, SI Appendix A).

**Fig. B1:**
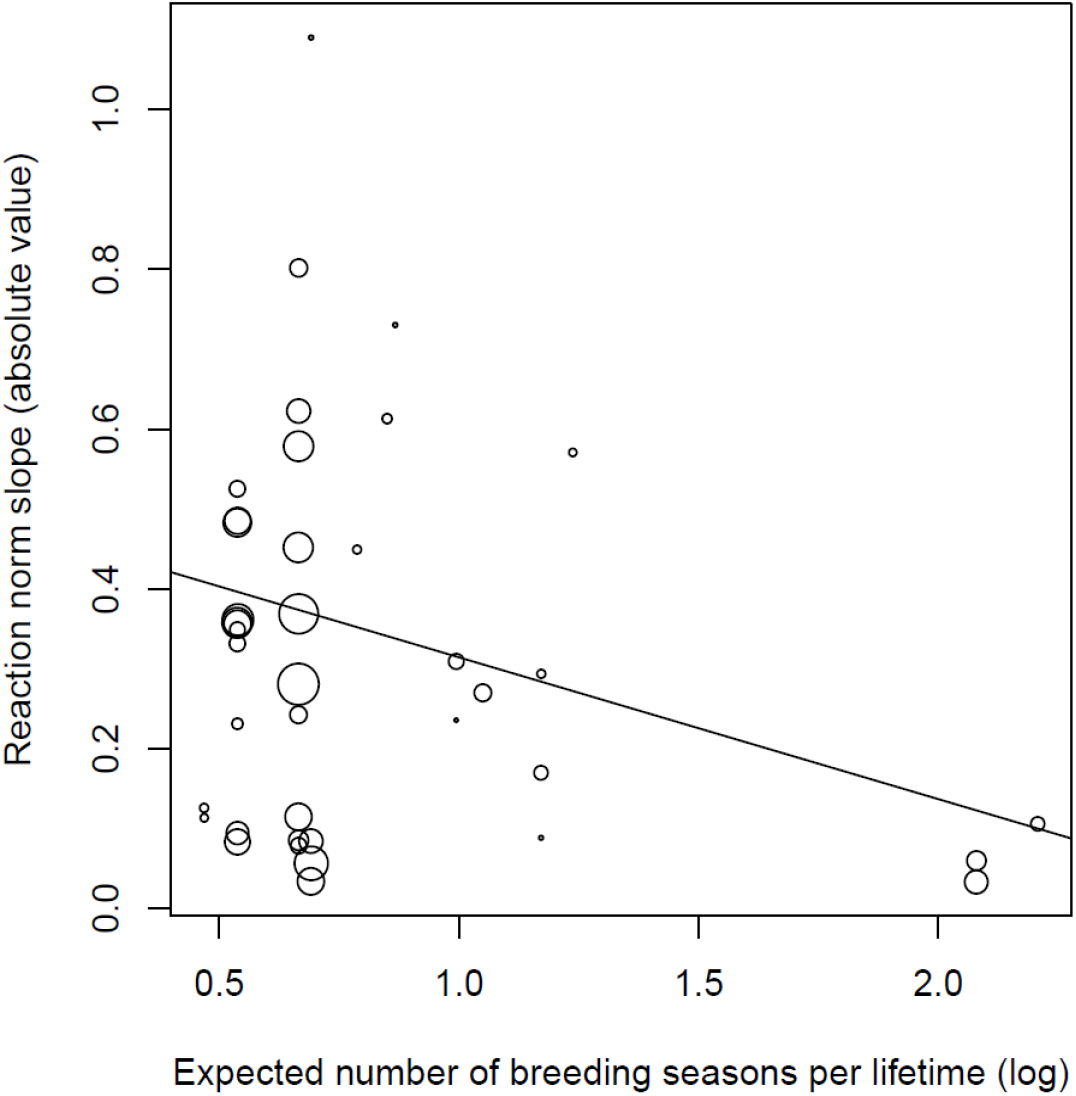
Absolute reaction norm slopes in response to temperature plotted as a function of the log number of breeding seasons a sexually mature individual expects to experience in its lifetime, for 23 populations of 13 species included in Radchuk et al. (2019)’s meta-analysis. Each point represents one reaction norm (predictor-response variable combination) slope estimate in a given population. Point size is inversely proportional to the standard error of the mean reaction norm slopes.

Although Radchuk et al. (2019) include year as a random effect in their meta-analysis, covariation with temperature variables due to directional climate warming across the duration of the study periods can make it difficult to be certain that the observed changes in lay-date are due to plasticity rather than microevolutionary change. This is a potential problem, because species with shorter generation times may be expected to evolve faster and track the moving optimum better via microevolution, and this is in the same direction as our prediction that bet-hedging should lead to adaptive increases in plasticity for species with shorter lifespans. Due to the low sample sizes, we are unable to restrict our analyses to only those studies that can estimate the relative roles of plasticity and microevolution (i.e. those with quantitative genetic analyses with multi-year individual-level data and population pedigrees). However, the majority of these more detailed studies are able to conclude that plasticity is the greater contributor to the observed response (Charmantier et al. 2008; Matthysen et al. 2011; Mihoub et al. 2012; Tarka et al. 2015; Thorley and Lord 2015), and this would support our contention that differences in plasticity rather than microevolution are driving the more rapid change in lay-dates in species with shorter lifespans.

Another caveat with this analysis it that it assumes that spring phenology correlates strongly with reproductive output for the entire season, but some populations here are able to re-nest later in the same season. This not only makes any plasticity or lack thereof in spring phenology (e.g. arrival date, first egg laying date) less important for yearly reproductive output, but also reduces the scope for bet hedging to act on spring phenology due to increasing opportunities for additive fitness accumulation. Interestingly, the data here for blue tits (*Cyanistes caeruleus*) and great tits (*Parus major*) include estimates for several populations, spanning a latitudinal gradient from northern to southern Europe, and populations at higher latitudes should experience much lower chances of re-nesting later within the same season. However, there was no significant difference in the magnitude of phenotypic plasticity spring phenology responses to temperature across latitudes, indicating that possibility of re-nesting may not be an important factor affecting fitness accumulation. One reason for this might be the strong seasonal decrease in reproductive success in these and other avian species, which reduces the impact of re-nesting attempts on any breeding success (Rowe et al. 1994; Verhulst et al. 1995; Verhulst and Nilsson 2008). However, it is clear that further studies are needed to resolve many of these issues properly.

Although we model the degree of investment in plasticity in terms of the slope of the reaction norm (*β*), which perhaps represents the most obvious source of any costs of plasticity, it should be noted that the results here should also hold for other types of increased investment in phenotype-environment matching with increasing costs and/or diminishing returns (Rago et al. 2019). For example, bet-hedging could also promote greater investment in the accuracy of plastic adjustments in the form of improved information gathering and cognitive processing, or in reductions in biological error during phenotypic expression (Westneat et al. 2015). Individuals that are able to adjust their phenotype faster, more often or more accurately might be expected to reduce variation in fitness outcomes by remaining as close as possible to the optimal phenotype and thus the peak fitness returns. However, as we explore in SI Appendix B, alternative formulations of costs of plasticity can change this prediction. If costs are paid not in terms of reaction norm slope itself (or not only), but rather in relation to the absolute amount of phenotypic change required each time (production costs), increased plasticity may in some cases lead to higher variance in fitness. This converse effect arising from an alternative costs of plasticity scenario could therefore cause conservative bet-hedging to favour a *less* plastic phenotype in systems where fitness accumulates multiplicatively (i.e. shallower reaction norm slopes evolve in low *n* scenarios in figs. S5B and S4B). Overall, whether genotype-level bet-hedging leads to an increase, or a decrease, in plasticity relative to individual-level optimization, the evolved degree of plasticity is expected to minimise fitness variances arising from costly phenotype-environment mismatches as well as from any costs of plasticity.

Previous theoretical treatments of the evolution of plasticity have often involved maximization of long-term geometric mean fitness, rather than just the short-term individual arithmetic mean fitness benefits. For example, in models of evolution of plasticity alongside bet-hedging (e.g. Botero et al. 2015; Tufto 2015), the bet-hedging benefits of plasticity are implicitly included when evolving the optimum degree of plasticity from the long-term genotype point-of-view. Consistent with our findings, Tufto’s (2015) scenarios with coarse spatial grain (without microenvironmental variability, Fig. 3 in Tufto 2015) consistently produced steeper reaction norm slopes than those with fine-grained environments (with microenvironmental variability, Figs. 2 and 4 in Tufto 2015). However, this trait of reaction norm slope co-evolves with the elevation of the reaction norm, and with random phenotypic noise (a diversifying bet-hedging effect, just termed ‘bet-hedging’), while tracking temporally autocorrelated macroenvironmental variability. Any change in the mean reaction norm slope is therefore seen in concert with simultaneous changes in these other modes of adaptation, and no part of it is interpreted as the conservative bet-hedging effect of adaptive reaction norms slopes that we isolate here.

Among published papers that have posited some bet-hedging effect leading to seemingly suboptimal effects on the degree of plasticity itself (Scheiner and Holt 2012; Scheiner 2013; King and Hadfield 2019), the results are only superficially similar to those presented here. These have identified the extreme outcomes of reaction norms steeper than 1 (‘hyperplasticity’) or negative reaction norms (‘negative plasticity’). However, all these models specifically consider developmental (irreversible) plasticity, and investigate whether phenotypes should respond to environmental conditions when the autocorrelation of the environment at time of development and at a single time of selection varies. Hyperplasticity, or negative plasticity, can then be favoured by specific (predictable) patterns of temporal autocorrelation either among or within generations, and especially if compensating for maladaptive changes in (mean) reaction norm intercept (fig. 4 in (King and Hadfield 2019)). Additionally, hyperplasticity (in response to developmental conditions) can arise as a diversifying bet-hedging strategy under certain ecological or life-history scenarios, such as when high juvenile dispersal (post-development but pre-selection) leads to related individuals experiencing a wide range of selective environments (Scheiner 2013). In contrast, we use a much simpler model and life-history, assuming that in the absence of plasticity the phenotypes will match mean environmental conditions. This allows us to isolate the effect of conservative bet-hedging lowering individual variance in fitness at the cost of lower expected individual fitness. It is difficult to detect whether such a bet-hedging effect is simultaneously at play in these and other complex models cited above, but it is especially likely in scenarios with low spatial variability and low dispersal.

To our knowledge, there have been no previous attempts to separate out the usual individual-level arithmetic mean fitness benefits of plasticity from the genotype-level bet-hedging advantages, as we have done here. In doing so, we have revealed a novel realization of conservative bet-hedging. Under one of the most common scenarios for costs of plasticity (scaled to the slope of the reaction norm), we find additional adaptive levels of plasticity due to plasticity’s effects in reducing individual fitness variance. This geometric mean fitness effect on plasticity has previously either been ignored or conflated with the usual arithmetic mean fitness benefits of adaptive plasticity. In addition, we have shown that any effect of bet-hedging on the degree of plasticity should depend upon the degree of multiplicative fitness accumulation, either due to coarser environmental grains (high *g*) and/or fewer time steps prior to reproduction (low *n*). It remains to be formally tested whether these predictions are upheld in empirical observations, for example in the exact degree of adaptive plasticity associated with fast versus slow life histories evolving under more or less variable environmental conditions (Ratikainen and Kokko 2019; e.g. Wright et al. 2019). However, we suggest here (Box 1) that such patterns may be visible in long-term datasets on avian breeding phenology in response to climate change (Radchuk et al. 2019). Similarly, we might expect to see these same types of among-species patterns in Debat & David’s (2001) assessments of the degree of physiological plasticity in terms of homeostatic ability to maintain a phenotype (e.g. body temperature, metabolic rate) irrespective of environmental variation. One important issue here remains the nature of any costs of plasticity (see SI Appendix B), and so our results suggest that we will gain a better understanding of the evolution of plasticity by partitioning out the different adaptive causes of plasticity on the short-term individual versus long-term genotypic levels.

## Acknowledgments

For discussions and comments on earlier versions of these ideas, we would like to thank Jarle Tufto, Luis-Miguel Chevin and Willem Frankenhuis. We thank Maja Tarka for discussions about the analysis of breeding phenology. TRH and IIR were supported by the Norwegian Research Council, grant 240008 to IIR on the Y oung Talented Researchers program, and grant 223257 to the Centre for Biodiversity Dynamics (CBD) at the Norwegian University of Science and Technology.

## Author contributions

TRH and IIR initiated the study. All authors developed the ideas in collaboration. TRH built the model, analysed the results and wrote the manuscript with input from all authors.

## Supporting Information

## Appendix A: Supplements to the main model

This part contains:

Figure S1: Simulation results from the model with discrete rather than overlapping generations.

Figure S2: Temporal (among-year) environmental variance as a function of spatial environmental grain.

Table S1: List of mathematical notation and parameter values.

Table S2: Results from alternative mixed effects models of phenological plasticity

**Figure S1:**
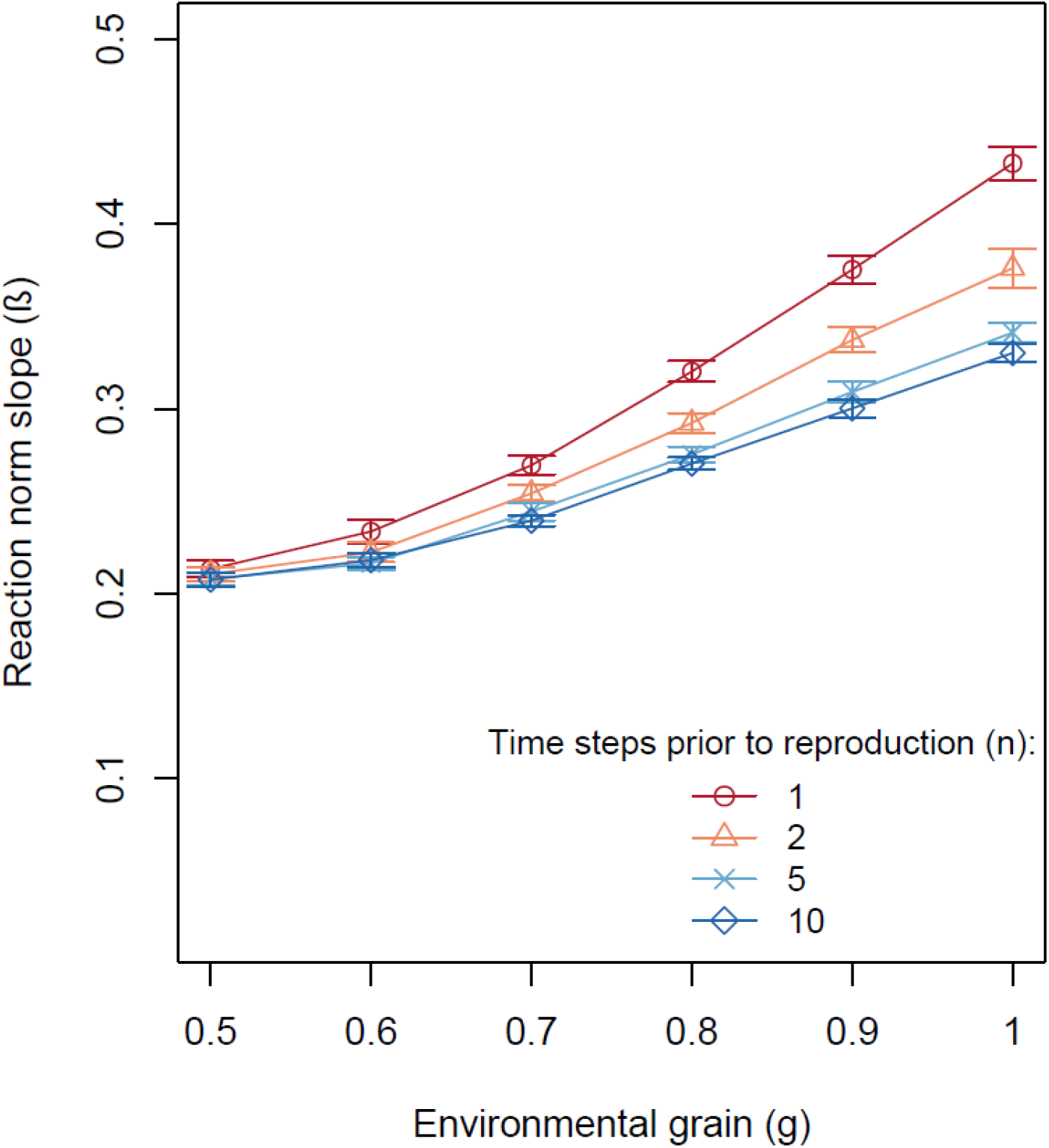
Mean evolved values of phenotypic plasticity *β* (reaction norm slope) from simulated scenarios with different values of environmental grain, *g* (x-axis), and number of time steps prior to reproduction, *n* (line colour, point type), when generations are discrete. Error bars show standard deviations in population means across 20 replicate runs. Baseline parameter values as shown in Table S1, except that *α* = 1 (100 % adult mortality between seasons).

**Figure S2:**
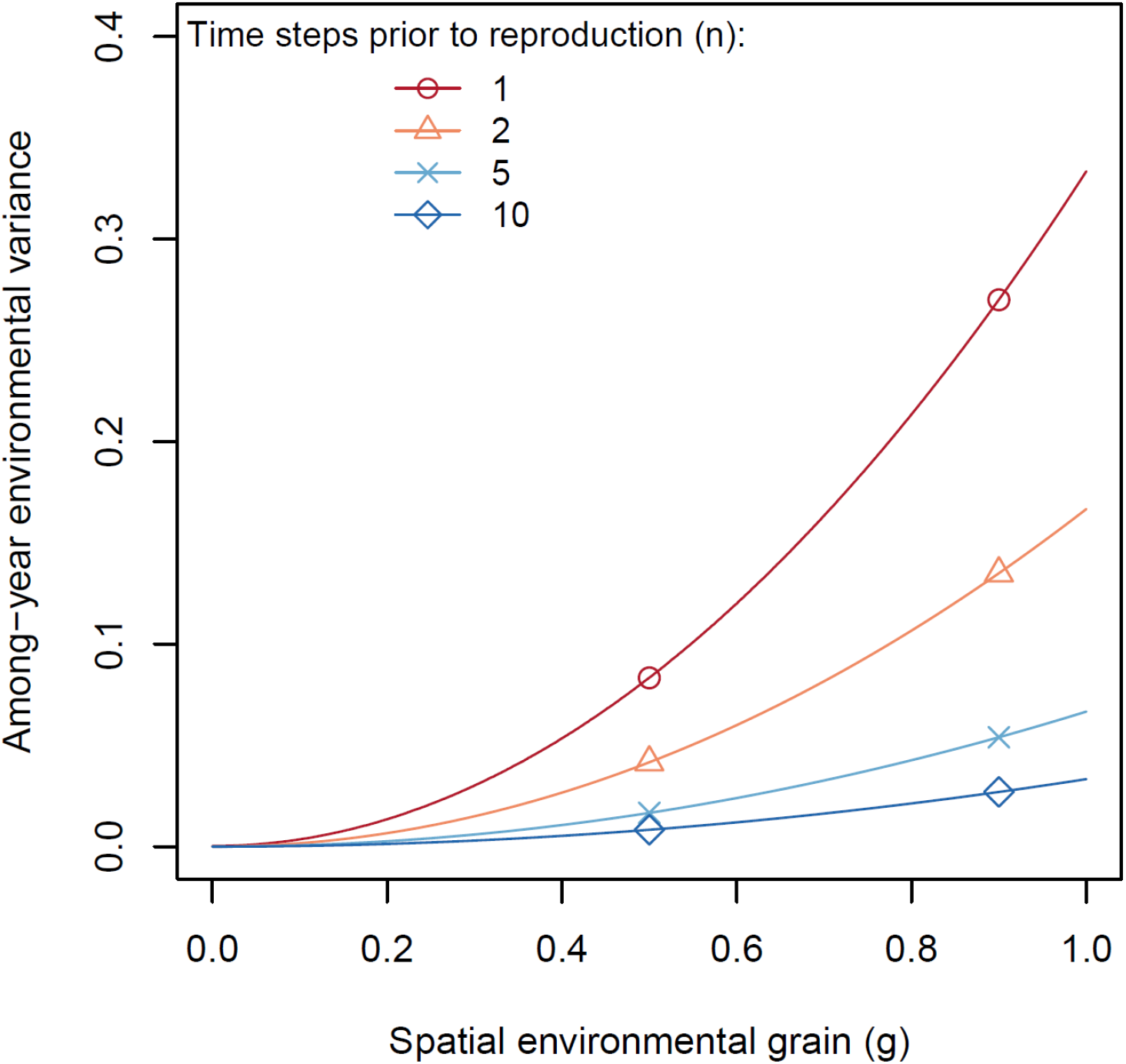
Among-year environmental variance for different numbers of time steps (each with their own, independently drawn environmental condition) within a year prior to reproduction (line colour, point type) and levels of spatial environmental grain (x-axis), representing the width of the distribution of possible environmental conditions in a given time step (see main text).

**Table S1:** List of mathematical notation used in the main text and their values in the results presented.

| Parameter or variable | Description | Range or baseline values |
| --- | --- | --- |
| $\beta_i$ | Gene for individual $i$'s reaction norm slope. | $0 \leq \beta \leq 1$ |
| $n$ | Number of time steps prior to reproduction | $n = \{1, 2, 5, 10\}$ |
| $E_t$ | Overall environmental condition in time step $t$ | $0 \leq E \leq 1$ |
| $e_{i,t}$ | Microenvironment experienced by individual $i$ in time step $t$ | $0 \leq e \leq 1$ |
| $g$ | Grain of environment: Similarity among $e_{i,t}$s at time $t$ | $g = \{0.4, 0.5, \dots, 1\}$ |
| $\alpha$ | Between-season mortality | Baseline 0.5. ($\alpha=1$ in Fig. S1) |
| $m$ | Probability of mutation | Baseline 0.005 |
| $m_\delta$ | Size of mutations (standard deviation of Gaussian distribution around parent's $\beta$ from which offspring's $\beta$ is drawn) | Baseline 0.05 |
| $K$ | Carrying capacity. | Baseline 5000 |
| $r$ | Intrinsic rate of increase (maximum reproductive success) | Baseline 2 |
| $\sigma$ | Standard deviation of the Gaussian function determining fitness decay as $z_{i,t}$ deviates from $e_{i,t}$ | Baseline 0.4 |
| $z_{i,t}$ | Phenotype of individual $i$ in time step $t$ | $0 \leq z_{i,t} \leq 1, z_{i,t} = \beta_i e_{i,t}$ |
| $W_i$ | Fitness (relative reproductive success) of individual $i$ | Eq. 1 |
| $s_1, s_2$ | Linear and exponential parameter for costs related to reaction norm slope (plasticity maintenance costs) | Baseline $s_1=1, s_2=2$ (varied in SI Appendix B) |
| $u_1, u_2$ | Linear and exponential parameter for costs related to phenotypic updating (plasticity production costs) | Baseline $u_1=0, u_2=0$ (varied in SI Appendix B) |

**Table S2:**
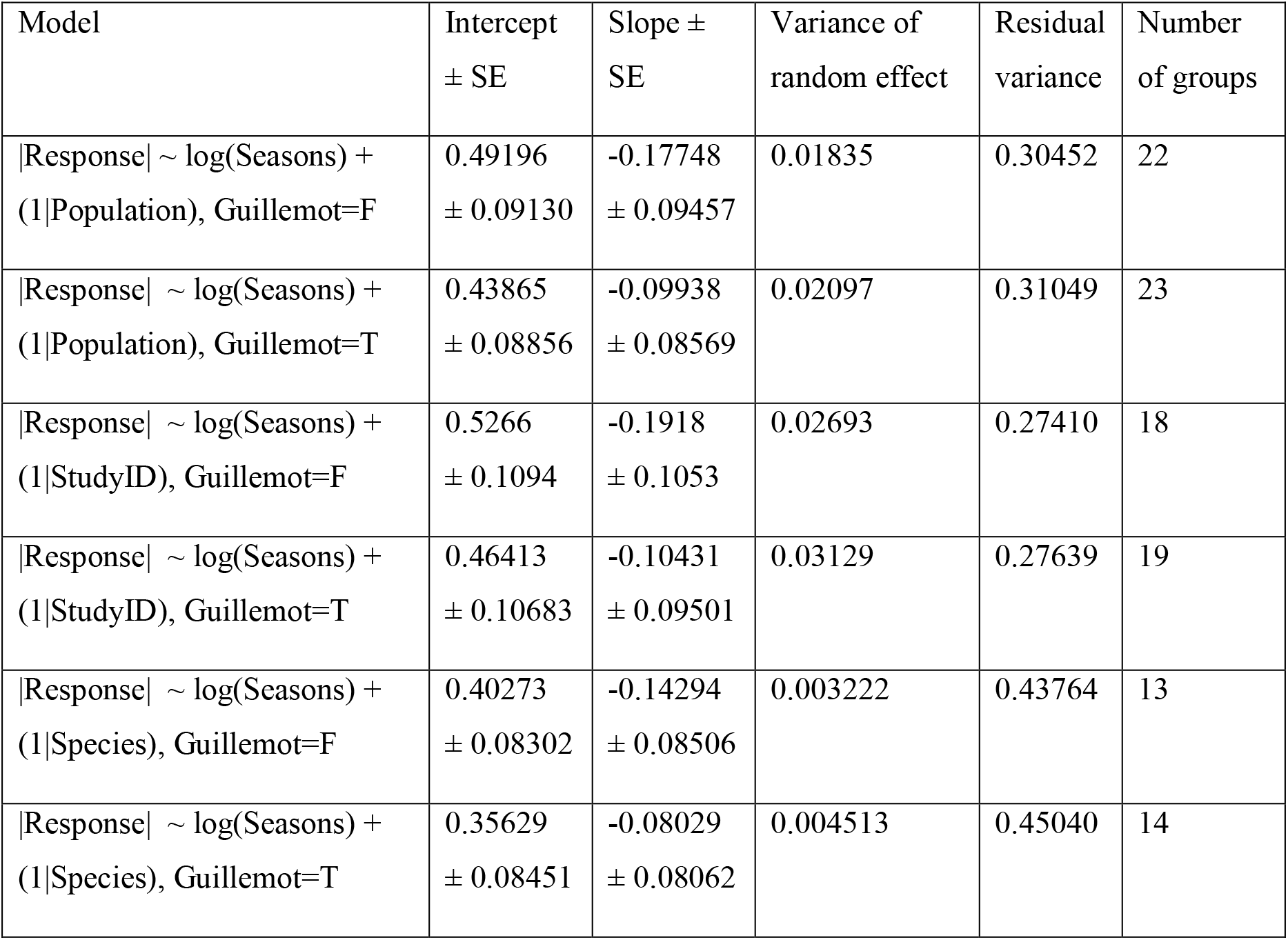
Results from alternative statistical models of the degree of plastic phenological (reaction norm ‘Slope’) responses to temperature. All models are linear mixed-effects models (lmer) with observation weights equal to 1/SE(Response), performed in the R version 3.6.1 (R Core Team 2019) using the package lme4. ‘Guillemot’: Include guillemot in analysis (T) or not (F). Number of groups indicates the levels within the random effect (‘Population’, ‘StudyID’ or ‘Species’). The first entry model is the one presented in the main text (Box 1, Fig. B1).

## Appendix B: Alternative costs of plasticity

Here we consider alternative formulations of the costs of plasticity, where costs do not (or not only) depend on the reaction norm slope directly, but possibly also on how much the phenotype is updated between consecutive time steps (production costs):

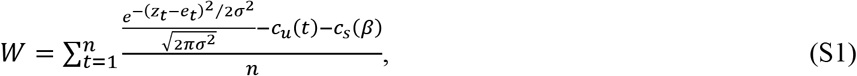

where *c*_s_(*β*) (*s* for “slope”) are the costs paid for simply ‘having’ a certain degree of plasticity (this function was *β*^2^ in the baseline model), and *c*_u_(*t*) (*u* for ‘update’) are the costs for updating the phenotype between time steps *t* and *t*-1. We here let *c*_s_(*β*) = *s*_1_ *β*^*s*_2_^, and *c*_u_(*t*) = *u*_1_ ļ(*z_t_* – *z*_*t*-1_)|*u*^2^, and typically set *s*_2_ = 1 and *u*_2_ = 1. The parameters *s*_1_ and *u*_1_, determine the steepness of the cost functions *c*_s_ and *c*_u_ in the linear case, whereas *s*_2_ and *u*_2_ can deviate from 1 to create non-linear cost functions.

When including *c*_u_, the question of *z*_0_ as the starting phenotype (or phenotype at ‘development’), becomes relevant – especially for scenarios with low *n* (number of time steps prior to reproduction). We let these phenotypes be dependent upon the environmental fluctuations in the same way as at the other times (i.e. *z*_0_ = *β*_*e*_0__), which also depends on the organism’s reaction norm slope. We also ensure that environmental grain affects *e*_0_ in the same way as it does the environments at other time steps, such that the starting phenotypes of individuals in coarse-grained environments and/or individuals with shallow reaction norm slopes are more similar than those of the individuals in fine-grained environments and/or with steeper reaction norm slopes.

### Predictions

We generate predictions for this model in the same way as for the model in the main text (Fig. 1; Model Setup). We are interested in the arithmetic and geometric mean *W* across all possible environments offered by different *β* values (reaction norm slopes). With the model in the main text considering only *c*_s_(*β*), we covered ‘all possible environments’ simply by generating 10^7^ random numbers from the same distribution as the environmental fluctuations used in the simulation model. However, now another aspect of environmental variation becomes important: ‘all possible *changes* in environment between two consecutive time steps’. Since the environmental fluctuations are uniformly distributed between −1 and 1, the absolute value of these changes has a triangular distribution with a lower bound and mode of 0, and an upper bound of 2. Similarly, the phenotypes expressed by an individual with reaction norm slope *β* will be uniformly distributed phenotypes between-*β* and *β*, and the individual will thus experience phenotypic changes that are triangle distributed with a lower bound and mode of 0, and an upper bound of *2β*. (Observe that if the environment shifts from the lowest to the highest value possible in back-to-back time steps from −1 to 1, or *vice versa*, the phenotype shifts from-*β* to *β*, a shift of magnitude 2*β*.) This distribution has a mean of 2*β*/3 and variance of 2*β*^2^/9. Since we unfortunately have been unable to fully identify the precise mean and variance of *W* for these scenarios, we chose to simply use the differences between the 10^7^ randomly generated numbers to approximate the ‘all possible changes in environment between two consecutive time steps’ over which to calculate fitness. This approximates the properties of the triangular distribution described above very closely (R^2^ = 0.95968, and simulating 10^8^ environments only increases this to 0.95997).

Note that we assume no temporal autocorrelation in the environment (each environmental value is i.i.d.). Whilst including temporal autocorrelation would indeed change the optimal reaction norm slopes, for example due to most changes between consecutive time steps becoming smaller, the effects of different reaction norm slopes in increasing or decreasing fitness variance would remain the same.

Fig. S3 and S4 show predictions for a variety of different cost scenarios including linear and quadratic *c*_Update_ and *c*_Slope_. Note that the geometric mean fitness functions in this figure are calculated using the approximation *W*_geom_ = E[*W*] − 2 Var[*W*] / E[*W*], since the exact solution exp{E[log(*W*)]} is not defined when *W* contains negative values, as occurs for many x-axis values in these scenarios.

**Figure S3:**
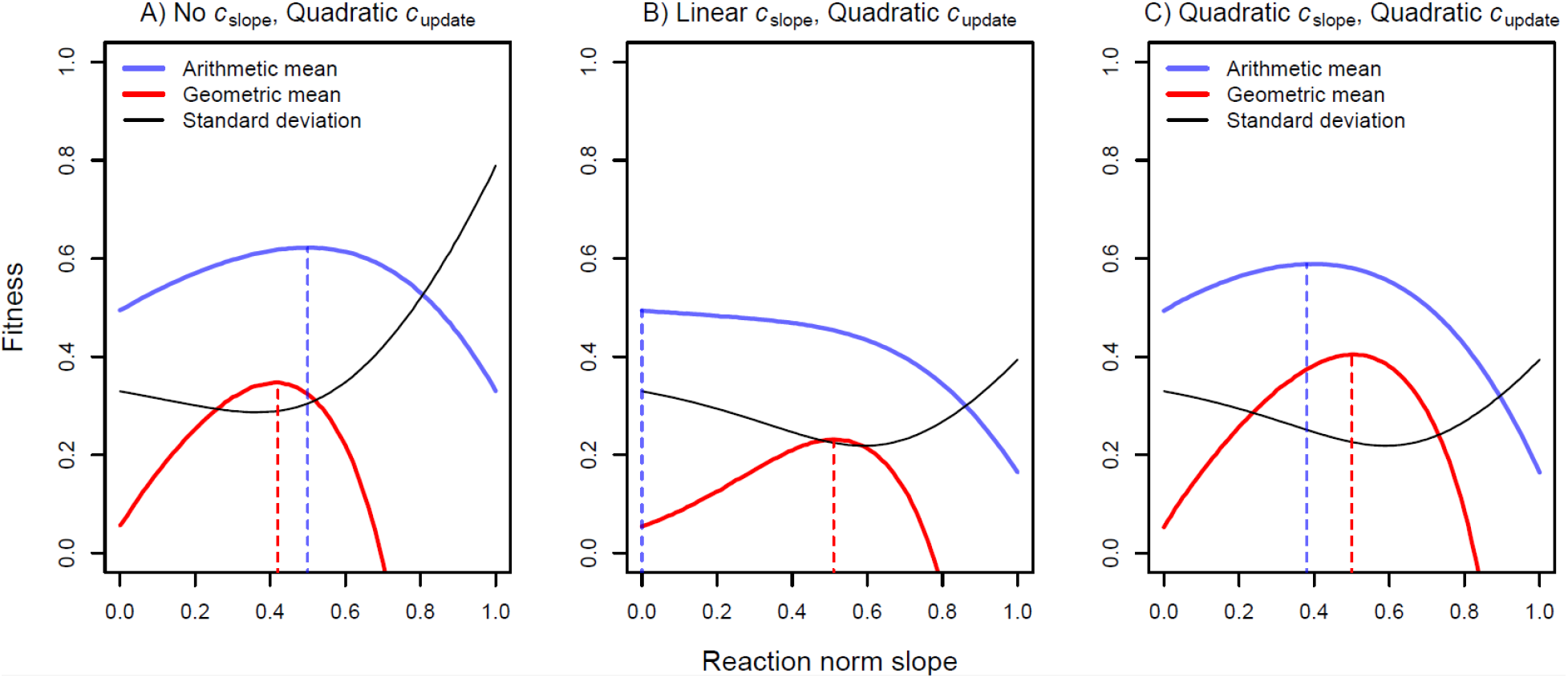
How plasticity affects fitness values for arithmetic mean (blue lines), geometric mean (red lines), and variance (black lines), for different variants of costs of plasticity. Dotted vertical lines show the maxima of the corresponding functions. A) Costs increase quadratically with how much the phenotype is updated between consecutive time steps – cost function coefficients are *s*_1_=0, *s*_2_=0, *u*_2_=1 and *u*_2_=2. B) Costs increase linearly with reaction norm slope, and quadratically with how much the phenotype is updated between time steps – cost function coefficients are *s*_1_=0.5, *s*_2_=1, *u*_2_=0.5 and *u*_2_=2. C) Costs increase quadratically with both reaction norm slope and phenotypic updating – cost function coefficients are *s*_1_=0.5, *s*_2_=2, *u*_2_=0.5 and *u*_2_=2. Since both types of costs are used in B and C, we chose less severe costs to avoid too low fitness.

**Figure S4:**
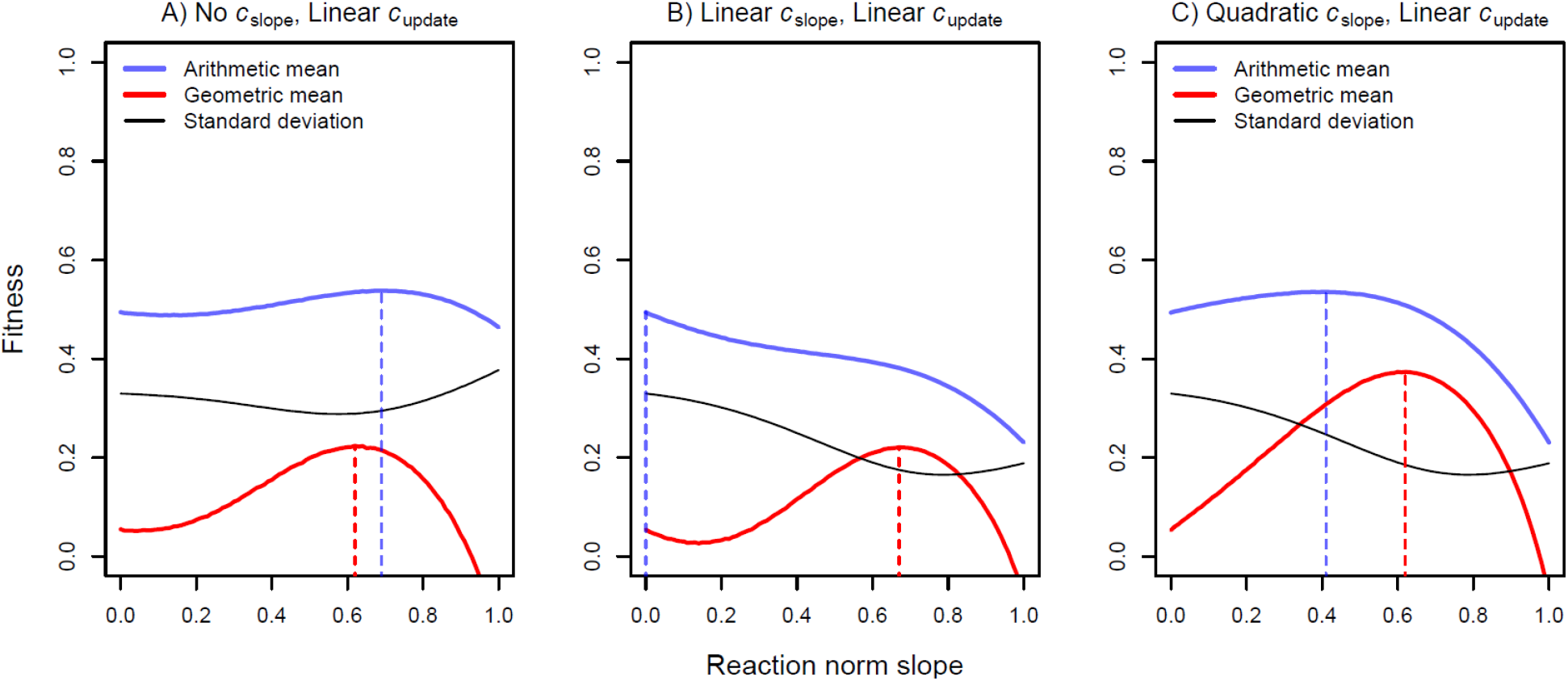
How plasticity affects fitness values for arithmetic mean (blue lines), geometric mean (red lines), and variance (black lines), for different variants of costs of plasticity. Dotted vertical lines show the maxima of the corresponding functions. A) Costs increase linearly with how much the phenotype is updated between consecutive time steps – cost function coefficients are *s*_1_=0, *s*_2_=0, *u*_2_=0.8 and *u*_2_=1. B) Costs increase linearly with reaction norm slope and phenotypic updating – cost function coefficients are *s*_1_=0.5, *s*_2_=1, *u*_2_=0.4 and *u*_2_=1. C) Costs increase quadratically with reaction norm slope and linearly with phenotypic updating – cost function coefficients are *s*_1_=0.5, *s*_2_=2, *u*_2_=0.4 and *u*_2_=1. Since both types of costs are used in B and C, we chose less severe costs to avoid too low fitness.

Note that when including *c*_Update_, fitness variance across environments is no longer strictly decreasing as *β* increases (black lines in Figs. S3 and S4 have an intermediate minimum). This is because although the variance in the fitness *gains* does decrease with *β* (eq. 1), the variance of the fitness costs (*c*_u_) increases (Var[*c*_u_]=0 only when *β*=0). This is in contrast with the effect of *c*_s_, which is a constant, and thus has Var[*c*s]=0 for all *β* (Fig. 1, main text). When this intermediate minimum occurs at lower *β* values than the *β* maximizing arithmetic mean fitness (as in Figs. S3A and S4A), the geometric mean fitness maximum is drawn towards this point, and bet-hedging is predicted to actually lead to shallower reaction norm slopes – not steeper, as in the result shown in the main text. This is counterintuitive given our argument in the main text, but makes sense under this particular ‘production cost’ structure. The degree of plasticity that maximizes arithmetic mean fitness across environments may in fact thus be ‘too’ plastic from a bet-hedging perspective. Because fitness becomes more variable as you pay the variable costs related to successive phenotypic changes, geometric mean fitness across generations may be optimized if individuals are less plastic than they would be if they were simply optimizing their arithmetic mean fitness. Matching the environmental optimum less well, on average, may therefore be favoured by bet-hedging if the fitness costs you need to pay in order to match the optimum are themselves prohibitively variable.

However, in Figs. S3B and S4B (with linear and quadratic *c*_s_, respectively) it is a non-plastic strategy that maximizes arithmetic mean fitness (blue line peaks at *β*=0), whereas geometric mean fitness favours a plastic strategy closer to the one that minimizes fitness variance. In Figs. S3C and S4C, geometric mean fitness similarly favours a more plastic strategy than does arithmetic mean fitness, as in the main text. Thus, the direction of the effect of conservative bet-hedging on reaction norm slopes may depend on the exact formulation of the costs of plasticity, as well as the benefits of accurately matching the phenotype to the environment (the shape of the fitness function), and these are likely to depend on the specific traits in question.

### Evolutionary simulations

To demonstrate the effect of conservative bet-hedging causing shallower reaction norm slopes (the opposite of the results in Fig. 1 and S1), we conducted evolutionary simulations as in the main text, exploring the range of the two parameters in our models that affect the degree of additive versus multiplicative fitness accumulation over time, which are environmental grain (*g*, correlations in environmental conditions among individuals at a given time) and the number of time steps *n* within a lifetime over which payoffs can accumulate prior to selection. With production costs of phenotypic changes but no maintenance costs of plasticity, we expect scenarios with low *n* and high *g* to produce less plastic phenotypes. Predictions and simulation results are shown in Figs. S5 (quadratically increasing costs, as in Fig. S3A) and S6 (linearly increasing costs, as in Fig. S4A), with all other model parameters as in Fig. 2 in the main text. The results presented in Fig. S5B confirm the predictions from Fig. S5A (and Fig. S3A), with reduced reaction norm slopes under conditions where bet-hedging is more important. However, in Fig. S6B we can see that although there is a slightly shallower reaction norm slope when there are fewer time steps before reproduction, there is a steeper reaction norm slope in coarser environmental grain. This is because as the amount of among-year environmental variation also decreases as environmental grain decreases (see main Results text and Fig. S2). Since environmental conditions are uniformly distributed between −1 and 1, it can be shown that among-year variance is equal to *g*^2^/3*n*, - i.e. increasing with higher *g* (coarser spatial grain) and lower *n* (fewer time steps per year with their own, independently drawn environmental conditions). Thus, this effect somewhat masks the trend of bet-hedging decreasing *β* in figures S5B and S6B (and amplifies it in Fig. 2 and S1), with bet-hedging “punishing” too steep reaction norms more strongly in Fig. S5 than in Fig. S6 (compare the slope of the fitness variance, black lines, in Fig. S5A versus Fig. S6A for *β* higher than the arithmetic mean fitness optima, dashed blue lines). A reasonable comparison to isolate the effect of additive versus multiplicative fitness accumulation from that of decreasing among-year variation could be to take the slope of a high *n* scenario (such as *n* = 10, or even higher) over different environmental grains as a baseline, and observe the relative effects of lowering *n*. As Fig. S2 also shows, these higher *n* scenarios experience much less change in among-year variation over *g* (blue lines are flatter in Fig. S2). Therefore, such a control reveals that for any given environmental grain, making fitness accumulation increasingly multiplicative favours shallower reaction norm slopes in Fig. S6B as well.

**Figure S5:**
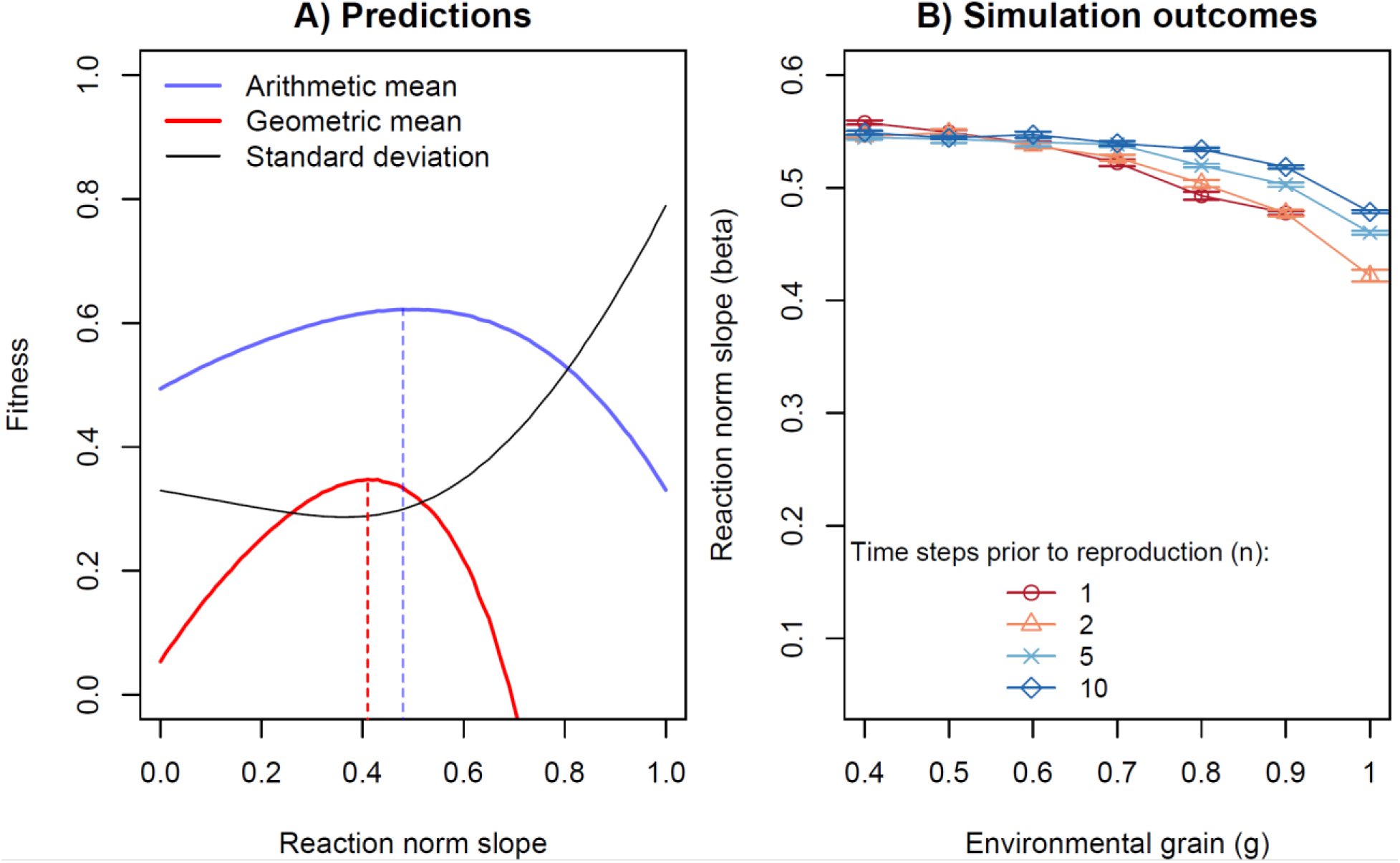
Comparison of predictions (A) and outcomes of evolutionary simulations (B) when the costs of plasticity increase quadratically with how much the phenotype is updated, but there is no cost to the reaction norm slope itself. Specifically, the cost function coefficients used are *s*_1_=0, *s*_2_=0, *u*_2_=1 and *u*_2_=2, as in Fig. S2A. Other model settings are as for the results presented in the main text, with parameter values shown in table S1. Error bars show standard deviations in mean population *β* across 20 replicate simulation runs. Note that no populations survived until the end of the simulations in runs where *n*=1 and *g*=1.

**Figure S6:**
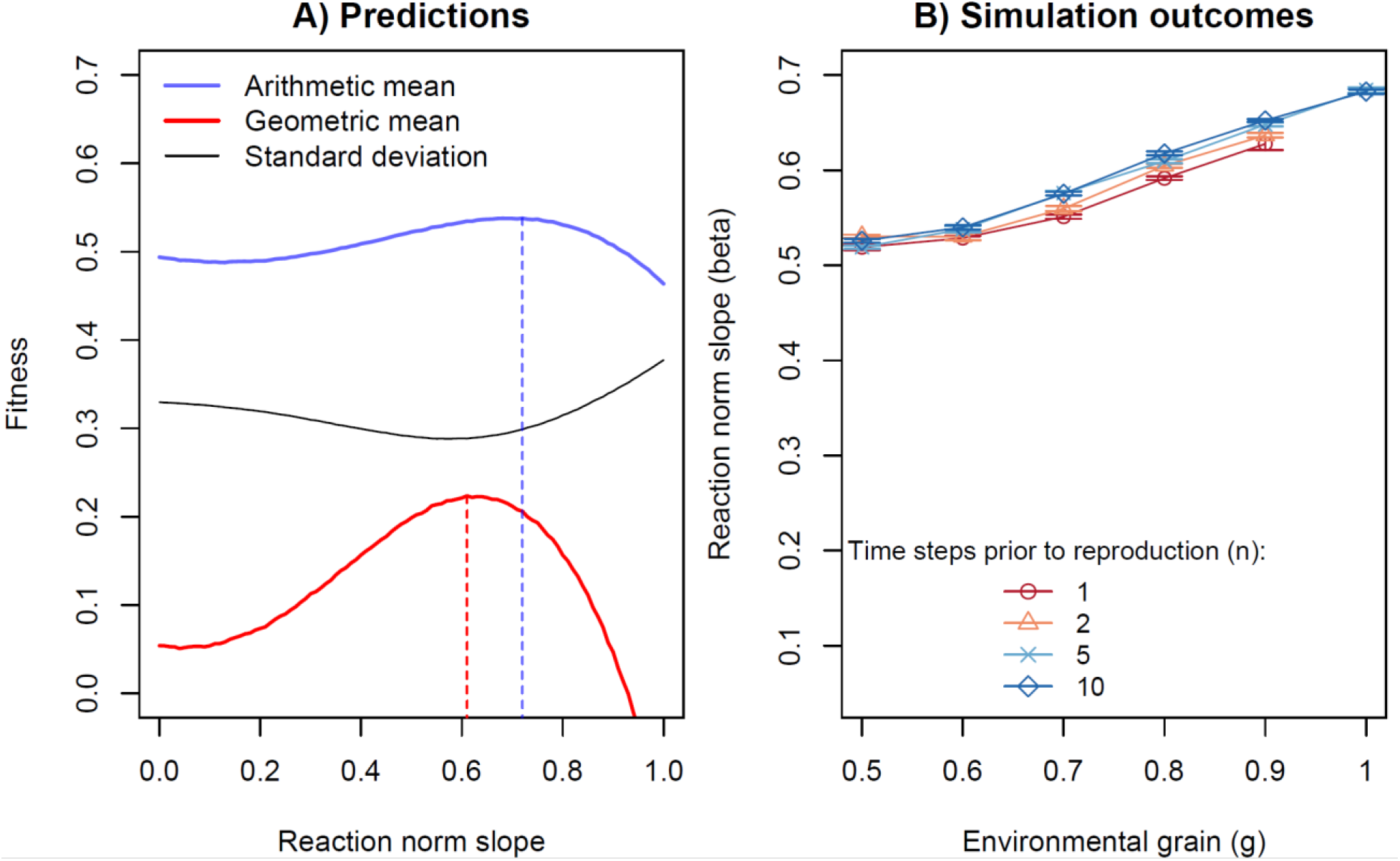
Comparison of predictions (A) and outcomes of evolutionary simulations (B) when the costs of plasticity increase linearly with how much the phenotype is updated, but there is no cost to the reaction norm slope itself. Specifically, the cost function coefficients used are s1=0, s2=0, u2=0.8 and u2=1, as in Fig. S3A. Other model settings are as for the results presented in the main text, with parameter values shown in table S1. Error bars show standard deviations in mean population β across 20 replicate simulation runs. Note that no populations survived until the end of the simulations in runs where n=1 or 2 and g=1.

## Appendix C: R code for the evolutionary simulations

~~~
rm(list=ls())
library(RColorBrewer)
##cost(): Calculate costs of plasticity
## Arguments:
## x: Reaction norm slope (for c_s) or magnitude of phenotypic update (for c_u)
## steepness: Slope and exponent parameters (s_1,s_2 or u_1,u_2)
cost <- function(x,steepness=c(1,1)){
return(steepness [1] *x^steepness[2])
}
## sim(): Individual-based simulation.
## Arguments:
## n: number of time steps/instances before reproduction, over which to gather resources.
## alpha: Between-year mortality
## m.rate: Mutation rate. Default 0.005
## m.size: Sd of the normal distribution around the old gene value from which new gene value is drawn.
## K: Carrying capacity. Default 5000. Density dependence is ‘ceiling’ type.
## T: Number of years/generations.
## reps: Number of replicate simulations. Default 1
## plot: Logical. Whether to plot population trajectories, or not (default not).
## grain_env: Correlation among individual in environments experienced in a time step. [0,1]
## r: Intrinsic rate of increase Default 2.
## costvar:How are costs of plasticity implemented? “slope” (only c_s, default), “update” (c_u) or “both”
## steepness: Cost function parameters s_1,s_2,u_1,u_2. See cost() function above.
## Title: Plot title. Default blank
## decayvar:How does fitness decay with mismatch? “gaussian” (default), “exponential” or “linear”
## decay: How steeply does fitness decay with mismatch? Only used if decayvar=“exponential”.
## Output: A list of two elements:
## [[1]] nstorage: Matrix of population sizes for each replicate (row) at each time step (column)
## [[2]] zstorage: Matrix of mean RN slope for each replicate (row) at each time step (column)
sim <- function(n,alpha=0.2,m.rate=0.005,m.size=0.05,K=5000,T=2000,reps=1,plot=FALSE,r=2, grain_env=0.1,costvar=“slope”,title=““,steepness=c(1,1,1,1),decayvar=“gaussian”,decay=5){
 if(plot){
  plot(1:T,rep(1,T),type=“n”,ylim=c(0,1),ylab=“Reaction norm slope”,xlab=“Year”,main=title)
 }
 nstorage <- zstorage <- matrix(NA,reps,T) #Storage matrices for population size and phenotypes.
 for(rep in 1:reps){
  #Initiate population
  N <- K #Starting pop. size is at carrying capacity
  pop <- runif(N) #Gene for reaction norm slope. Initialize uniform.
  for(Time in 1:T){
   #Record population traits
   zstorage[rep,Time] <- mean(abs(pop)) #Record plasticities
   nstorage[rep,Time] <- N #Record population size.
   W <- rep(0,N)
   # Developmental environment determining starting phenotypes (only necessary if including updatecosts)
   if(costvar==“update”||costvar==“both”){
    env <- runif(1)
    lower <- grain_env*env
    upper <- env + (1-grain_env)*(1-env)
    envs <- runif(N,lower,upper)*2-1 #Environments are between −1 and 1
    phens <- pop*envs # Phenotype is Environment*RN Slope. RN Slope==1 ->
 phenotype==environment
   }
   for(Step in 1:n){
    if(costvar==“update”||costvar==“both”){prev.phens <- phens} # Record last time’s phenotype (only necessary if including updatecosts)
    this.W <- rep(0,N)
    #Find environment(s)
    env <- runif(1)
    lower <- grain_env*env
    upper <- env + (1-grain_env)*(1-env)
    envs <- runif(N,lower,upper)*2-1 #Environments are between −1 and 1
    #Find individual phenotypes
    phens <- pop*envs # Phenotype is Environment*RN Slope. RN Slope==1 ->
 phenotype==environment
    #Calculate fitnesses dependent on phenotype-environment mismatch (states-conds)
    this.W <- switch(decayvar,
        “exponential”= dexp(abs(phens-envs),decay)/decay,
        “linear”= 1-abs(phens-envs),
        “gaussian”= dnorm(phens-envs,0,0.4))
    W <- W + this.W #Sum fitnesses accumulated
    if(costvar==“update”||costvar==“both”){ #If cost depends on how much the phenotype changed
     W <- W - cost(abs(phens-prev.phens),steepness=steepness[3:4])
    }else if(costvar==“slope”||costvar==“both”){ #If cost depends on plasticity gene
     W <- W - cost(abs(pop),steepness=steepness[1:2])
    }
   }
   W <- r*(W/n) # Standardize fitness on n, multiply by intrinsic rate of increase W[W<0] <- 0
   if(round(sum(W)==0)){
    print(c(“Population extinct at time”,Time))
    break
   }
   #Selection
   if(alpha==1){ # If discrete generations (all adults die)
    NextN <- ifelse(sum(W)>K,K,round(sum(W))) # Find population size next year
    pop <- sample(pop,size=NextN,prob=W,replace=TRUE) # Choose offspring according to parent’s
fitness
   if(costvar==“update”||costvar==“both”) phens <- rep(0,NextN) # All offspring are adapted to ‘mean’ environmental conditions.
 #Mutation
 mut <- which(runif(NextN)<m.rate) #Select mutated individuals
 pop[mut] <- rnorm(length(mut),pop[mut],m.size)
}
else{ # If overlapping generations (some adults survive)
 Alive <- runif(N)>alpha # Random survival
 Alive <- which(Alive==TRUE) # Alive is a sequence of numbers of the individuals who survived.
 #How many ‘slots’ should be filled (i.e. how many offspring should be produced?)
 Offspring <- min(round(sum(W)),K-length(Alive))
 #Reproduction. IDs of those who get offspring.
 if(sum(W[Alive])>0){ #In this scenario, offspring die if no surviving parents.
  IDs <- sample(Alive,size=Offspring,prob=W[Alive],replace=TRUE)
 } else {
  print(c(“Extinct at time”,Time))
  break
}
 #Next generation is a combination of new and old individuals.
 pop <- c(pop[IDs],pop[Alive]) #IDs has length Offspring, so the Offspring first entries here are
new.
 if(costvar[1]==“update”) phens <- c(rep(0,Offspring),phens[Alive]) # All offspring are adapted to ‘mean’ env. conditions.
 #Mutation: Some of the offspring mutate
 mut <- which(runif(Offspring)<m.rate)
 pop[mut] <- rnorm(length(mut),pop[mut],m.size)
 NextN <- length(pop)
 }
 N <- NextN
}
 print(rep) # Ticker
 if(plot){
    lines(1:T,zstorage[rep,],col=rgb(1,0,0,0.1))
    lines(1:T,nstorage[rep,]/K,col=rgb(0,0,1,0.1))
   }
  }
 return(list(zstorage,nstorage))
}
#### Run scenarios ####
gs <- seq(0.4,1,by=0.1) #Environmental grain scenarios
ns <- c(1,2,5,10) #Number of time steps prior to reproduction
T <- 2000
K <- 5000
#Initialize storage
meanphens <- sdphens <- n <- sdn <- numeric(length(ns))
out <- array(0,dim=c(length(ns),4,length(gs)))
dimnames(out) <- list(ns,c(“meanphens”,”sdphens”,”n”,”sdn”),gs)
# Not hard-coded - change the parameters you want in function call
for(g in gs){
 for(i in ns){
  #Run simulation
  test <-
sim(n=i,alpha=0.5,reps=20,plot=TRUE,grain_env=g,decayvar=“gaussian”,costvar=c(“update”,”linear”),st eepness=1)
  #Record summary statistics
  repmeans <- apply(test[[1]][,(T-1000):T],2,mean,na.rm=TRUE) #Phenotypes in the last 1000 generations.
  meanphens[which(i==ns)] <- mean(repmeans,na.rm=TRUE) #Mean phenotypes across replicates
  sdphens[which(i==ns)] <- sd(repmeans,na.rm=TRUE) #Sd phenotypes across replicates
  rep_mean_n <- apply(test[[2]][,(T-1000):T],1,mean,na.rm=TRUE) #Population sizes
  n[which(i==ns)] <- mean(rep_mean_n,na.rm=TRUE)/K #Mean population size across replicates
 (Relative to K)
  sdn[which(i==ns)] <- sd(rep_mean_n,na.rm=TRUE) #Sd population size across replicates
  print(c(g,i)) # Ticker
 }
 out[,,which(g==gs)] <- c(meanphens,sdphens,n,sdn)
}
saveRDS(out,file=“Your directory and file name.Rdata”) # Save output
#### Create results plots (Fig. 2 and S1) ####
#Read output
data <- readRDS(file=“Your directory and file name.Rdata”)
# Set diverging red-blue color scheme
pal <- brewer.pal(length(data[,1,1])+2,”RdBu”)
pal <- c(head(pal,length(pal)/2-1),tail(pal,length(pal)/2-1))
pchs <- c(1,2,4,5) # Point types for plotting - length(pchs) should be equal to length (ns)
plot(1:length(gs),data[1,1,1:length(gs)],ylim=c(0,0.9),type=“n”,xaxt=“n”,xlab=“Environmental grain
(g)”,ylab=“Reaction norm slope (beta)”)
legend(“bottomleft”,legend=ns,col=pal,lty=1,title=“Time steps prior to reproduction
(n): “,bty=“n”,pch=pchs,cex=0.9,pt.cex= 1.1)
axis(side= 1,at= 1:length(gs),label=gs)
for(i in 1:length(ns)){
 points(1:length(gs),data[i,1,1:length(gs)],pch=pchs[i],cex=1,col=pal[i])
 lines(1:length(gs),data[i,1,1:length(gs)],col=pal[i])
 for(j in 1:length(gs)){
  arrows(j,data[i,1,j],j,data[i,1,j]+data[i,2,j],angle=90,length=0.1,col=pal[i])
  arrows(j,data[i,1,j],j,data[i,1,j]-data[i,2,j],angle=90,length=0.1,col=pal[i])
 }
}
#### Calculate variance in fitness ####
## Gaussian W(m)
# b is the different plasticity slopes (should be a vector, e.g. seq(0,1,by=0.01))
# cost is whether costs depend on RN slope (“slope”, default) or how much the phenotype is updated
(“update”) or “both”
# steepness[l:2] determines linear and exponent of slopecosts (if cost= “slope” or “both”) - default 1, 2
# steepness[3:4] determines linear and exponent of updatecosts (if cost= “update” or “both”) - default 0.5,
1
# Returns a vector of:
# [1] Variance in fitness across environments
# [2] Arithmetic mean fitness across environments
# [3] Geometric mean fitness across environments (approximation)
# [4] Geometric mean fitness across environments (exact - but doesn’t work if some fitnesses are negative)
sim_gaus <- function(b,cost=c(“slope”),steepness=c(1,2,0.5,1)){
 es <- runif(1000000,-1,1) # Simulate random environments
 zs <- b*es # Phenotypes produced in each env.
 ws <- dnorm(tail(abs(zs-es),length(zs)-1),0,0.4) # Fitness benefits - first env. is “developmental env.” (in case of using updatecosts)
 slopecost <- cost(b,steepness=steepness[1:2]) # c_s
 updatecost <- cost(abs(head(zs,length(zs)-1)-tail(zs,length(zs)-1)),steepness=steepness[3:4]) # c_u
  if(cost==“slope”) {
   ws <- ws-slopecost
  } else if(cost==“update”){
   ws <- ws-updatecost
  } else if(cost==“both”){
   ws <- ws-slopecost-updatecost
 }
 logws <- log(ws)
 logws[which(is.nan(logws)==TRUE)] <- −10000
 return(c(var(ws),mean(ws),mean(ws)-(2*var(ws))/mean(ws),exp(mean(logws))))
}
#### Create predictions figures (S3 and S4) ####
bs <- seq(0,1,by=0.01) #For any RN slope
vars <- ameans <- gmeansapprox <- gmeansexact <- numeric(length(bs))
for(i in 1:length(bs)){
 tmp <- sim_gaus(bs[i],cost=“update”,steepness=c(0,0,1,2))
 vars[i] <- tmp[1]
 ameans[i] <- tmp [2]
 gmeansapprox[i] <- tmp[3]
 gmeansexact[i] <- tmp[4]
 print(i)
}
plot(bs,gmeansapprox,main=expression(“A) No “*italic(c)[slope]*”, Quadratic
“*italic(c)[update]),type=“l”,col=“Red”,xlab=““,ylim=c(0,1),ylab=“Fitness”,cex.lab=1.3,lwd=2)
lines(bs,ameans,lwd=2,col=rgb(0,0,1,0.6))
lines(bs,sqrt(vars))
arrows(bs[which.max(ameans)],-
0.05,bs[which.max(ameans)],max(ameans),length=0,col=rgb(0,0,1,0.6),lty=2)
arrows(bs[which.max(gmeansapprox)],-
0.05,bs[which.max(gmeansapprox)],max(gmeansapprox),length=0,col=“ Red”,lty=2)
legend(“topleft”,legend=c(“Arithmetic mean”,”Geometric mean”,”Standard deviation”), lwd=c(2,2,1),lty=c(1,1,1),col=c(rgb(0,0,1,0.6),”Red”,”Black”),bty=“n”)
~~~

## Literature cited

Auld, J. R., A. A. Agrawal, and R. A. Relyea. 2010. Re-evaluating the costs and limits of adaptive phenotypic plasticity. Proc. R. Soc. B Biol. Sci. 277:503–511.

Bonamour, S., L. M. Chevin, A. Charmantier, and C. Teplitsky. 2019. Phenotypic plasticity in response to climate change: The importance of cue variation. Philos. Trans. R. Soc. B Biol. Sci. 374.

Botero, C. A., F. J. Weissing, J. Wright, and D. R. Rubenstein. 2015. Evolutionary tipping points in the capacity to adapt to environmental change. Proc. Natl. Acad. Sci. 112:184–189.

Charmantier, A., R. H. McCleery, L. R. Cole, C. Perrins, L. E. B. Kruuk, and B. C. Sheldon. 2008. Adaptive phenotypic plasticity in response to climate change in a wild bird population. Science 320:800–803.

Chevin, L.-M., R. Lande, and G. M. Mace. 2010. Adaptation, plasticity, and extinction in a changing environment: towards a predictive theory. PLoS Biol. 8:e1000357.

Clauss, M. J., and D. L. Venable. 2000. Seed germination in desert annuals: An empirical test of adaptive bet hedging. Am. Nat. 155:168–186.

Cohen, D. 1966. Optimizing reproduction in a randomly varying environment. J. Theor. Biol. 12:119–129.

Crowley, P. H., S. M. Ehlman, E. Korn, and A. Sih. 2016. Dealing with stochastic environmental variation in space and time: bet hedging by generalist, specialist, and diversified strategies. Theor. Ecol. 9:149–161.

Debat, V., and P. David. 2001. Mapping phenotypes: Canalization, plasticity and developmental stability. Trends Ecol. Evol. 16:555–561.

Dempster, E. R. 1955. Maintenance of genetic heterogeneity. Cold Spring Harb. Symp. Quant. Biol. 20:25–32.

DeWitt, T. J., A. Sih, and D. S. Wilson. 1998. Cost and limits of phenotypic plasticity. Trends Ecol. Evol. 13:77–81.

Dingemanse, N. J., A. J. N. Kazem, D. Réale, and J. Wright. 2010. Behavioural reaction norms: animal personality meets individual plasticity. Trends Ecol. Evol. 25:81–89.

Donaldson-Matasci, M. C., C. T. Bergstrom, and M. Lachmann. 2013. When unreliable cues are good enough. Am. Nat. 182:313–27.

Elner, R. W., and R. Hughes. 1978. Energy maximization in the diet of the shore crab, Carcinus maenas. J. Anim. Ecol. 47:103–116.

Fox, R. J., J. M. Donelson, C. Schunter, T. Ravasi, and J. D. Gaitán-Espitia. 2019. Beyond buying time: The role of plasticity in phenotypic adaptation to rapid environmental change. Philos. Trans. R. Soc. B Biol. Sci. 374.

Franch-Gras, L., E. M. García-Roger, M. Serra, and M. J. Carmona. 2017. Adaptation in response to environmental unpredictability. Proc. R. Soc. B Biol. Sci. 284.

Frank, S. A., and M. Slatkin. 1990. Evolution in a variable environment. Am. Nat. 136:244–260.

Frankenhuis, W. E., K. Panchanathan, and H. Clark Barrett. 2013. Bridging developmental systems theory and evolutionary psychology using dynamic optimization. Dev. Sci. 16:584–598.

Furness, A. I., K. Lee, and D. N. Reznick. 2015. Adaptation in a variable environment: Phenotypic plasticity and bet-hedging during egg diapause and hatching in an annual killifish. Evolution 69:1461–1475.

Gerber, N., and H. Kokko. 2018. Abandoning the ship using sex, dispersal or dormancy: multiple escape routes from challenging conditions. Philos. Trans. R. Soc. Lond. B. Biol. Sci. 373.

Gillespie, J. H. 1974. Natural selection for within-generation variance in offspring number. Genetics 76:601–6.

Grantham, M. E., C. J. Antonio, B. R. O’Neil, Y. X. Zhan, and J. A. Brisson. 2016. A case for a joint strategy of diversified bet hedging and plasticity in the pea aphid wing polyphenism. Biol. Lett. 12:20160654.

Gremer, J. R., E. E. Crone, and P. Lesica. 2012. Are dormant plants hedging their bets? Demographic consequences of prolonged dormancy in variable environments. Am. Nat. 179:315–327.

Gremer, J. R., and D. L. Venable. 2014. Bet hedging in desert winter annual plants: Optimal germination strategies in a variable environment. Ecol. Lett. 17:380–387.

Haaland, T. R., and C. A. Botero. 2019. Alternative responses to rare selection events are differentially vulnerable to changes in the frequency, scope, and intensity of environmental extremes. Ecol. Evol. 9:11752–11761.

Haaland, T. R., J. Wright, and I. I. Ratikainen. 2019a. Bet-hedging across generations can affect the evolution of variance-sensitive strategies within generations. Proc. R. Soc. B Biol. Sci. 286:20192070.

Haaland, T. R., J. Wright, and I. I. Ratikainen. 2020. Generalists versus specialists in fluctuating environments: a bet-hedging perspective. Oikos 1–12.

Haaland, T. R., J. Wright, J. Tufto, and I. I. Ratikainen. 2019b. Short-term insurance versus long-term bet-hedging strategies as adaptations to variable environments. Evolution 73:145–157.

Hansen, T. F. 2018. Fitness in evolutionary biology. Preprints.org 1–26.

King, J. G., and J. D. Hadfield. 2019. The evolution of phenotypic plasticity when environments fluctuate in time and space. Evol. Lett. 395137.

Levins, R. 1962. Theory of fitness in a heterogeneous environment. I. The fitness set and adaptive function. Am. Nat. 96:361–373.

Lewontin, R. C., and D. Cohen. 1969. On population growth in a randomly varying environment. Proc. Natl. Acad. Sci. 62:1056–1060.

Lynch, M., and W. Gabriel. 1987. Environmental tolerance. Am. Nat. 129:283–303.

Matthysen, E., F. Adriaensen, and A. A. Dhondt. 2011. Multiple responses to increasing spring temperatures in the breeding cycle of blue and great tits (*Cyanistes caeruleus*, Parus major). Glob. Chang. Biol. 17:1–16.

McNamara, J. M. 1998. Phenotypic plasticity in fluctuating environments: consequences of the lack of individual optimization. Behav. Ecol. 9:642–648.

Merilä, J., and A. P. Hendry. 2014. Climate change, adaptation, and phenotypic plasticity: the problem and the evidence. Evol. Appl. 7:1–14.

Mihoub, J. B., N. G. Mouawad, P. Pilard, F. Jiguet, M. Low, and C. Teplitsky. 2012. Impact of temperature on the breeding performance and selection patterns in lesser kestrels Falco naumanni. J. Avian Biol. 43:472–480.

Parker, G. A., and J. Maynard Smith. 1990. Optimality theory in evolutionary biology. Nature 348:27–33.

Philippi, T., and J. Seger. 1989. Hedging one’s evolutionary bets, revisited. Trends Ecol. Evol. 4:41–44.

Plard, F., J. M. Gaillard, T. Coulson, A. J. M. Hewison, D. Delorme, C. Warnant, and C. Bonenfant. 2014. Mismatch between birth date and vegetation phenology slows the demography of roe deer. PLoS Biol. 12:1–8.

Poethke, H. J., T. Hovestadt, and O. Mitesser. 2016. The evolution of optimal emergence times: bet hedging and the quest for an ideal free temporal distribution of individuals. Oikos 125:1647–1656.

R Core Team. 2019. R: A language and environment for statistical computing. R Foundation for Statistical Computing, Vienna, Austria.

Radchuk, V., T. Reed, C. Teplitsky, M. van de Pol, A. Charmantier, C. Hassall, P. Adamík, F. Adriaensen, M. P. Ahola, P. Arcese, J. Miguel Avilés, J. Balbontin, K. S. Berg, A. Borras, S. Burthe, J. Clobert, N. Dehnhard, F. de Lope, A. A. Dhondt, N. J. Dingemanse, H. Doi, T. Eeva, J. Fickel, I. Filella, F. Fossøy, A. E. Goodenough, S. J. G. Hall, B. Hansson, M. Harris, D. Hasselquist, T. Hickler, J. Joshi, H. Kharouba, J. G. Martínez, J.-B. Mihoub, J. A. Mills, M. Molina-Morales, A. Moksnes, A. Ozgul, D. Parejo, P. Pilard, M. Poisbleau, F. Rousset, M.-O. Rödel, D. Scott, J. C. Senar, C. Stefanescu, B. G. Stokke, T. Kusano, M. Tarka, C. E. Tarwater, K. Thonicke, J. Thorley, A. Wilting, P. Tryjanowski, J. Merilä, B. C. Sheldon, A. Pape Møller, E. Matthysen, F. Janzen, F. S. Dobson, M. E. Visser, S. R. Beissinger, A. Courtiol, and S. Kramer-Schadt. 2019. Adaptive responses of animals to climate change are most likely insufficient. Nat. Commun. 10:3109.

Rago, A., K. Kouvaris, T. Uller, and R. Watson. 2019. How adaptive plasticity evolves when selected against. PLoS Comput. Biol. 15:1–20.

Ratikainen, I. I., and H. Kokko. 2019. The coevolution of lifespan and reversible plasticity. Nat. Commun. 10:538. Springer US.

Reed, T. E., S. W. Robin, D. E. Schindler, J. J. Hard, and M. T. Kinnison. 2010. Phenotypic plasticity and population viability: The importance of environmental predictability. Proc. R. Soc. B Biol. Sci. 277:3391–3400.

Reed, T. E., P. Warzybok, A. J. Wilson, R. W. Bradley, S. Wanless, and W. J. Sydeman. 2009. Timing is everything: Flexible phenology and shifting selection in a colonial seabird. J. Anim. Ecol. 78:376–387.

Rowe, L., D. Ludwig, and D. Schluter. 1994. Time, condition, and the seasonal decline of avian clutch size. Am. Nat. 143:698–722.

Scheiner, S. M. 2013. The genetics of phenotypic plasticity: XII: Temporal and spatial heterogeneity. Ecol. Evol. 3:4596–4609.

Scheiner, S. M., and R. D. Holt. 2012. The genetics of phenotypic plasticity. X. Variation versus uncertainty. Ecol. Evol. 2:751–767.

Seger, J., and H. J. Brockmann. 1987. What is bet-hedging? Oxford Surv. Evol. Biol. 4:182–211.

Simons, A. M. 2011. Modes of response to environmental change and the elusive empirical evidence for bet hedging. Proc. R. Soc. B Biol. Sci. 278:1601–1609.

Simons, A. M. 2014. Playing smart vs. playing safe: The joint expression of phenotypic plasticity and potential bet hedging across and within thermal environments. J. Evol. Biol. 27:1047–1056.

Simons, A. M., and M. O. Johnston. 2003. Suboptimal timing of reproduction in *Lobelia inflata* may be a conservative bet-hedging strategy. J. Evol. Biol. 16:233–243.

Slatkin, M. 1974. Hedging one’s evolutionary bets. Nature 250:704–705.

Starrfelt, J., and H. Kokko. 2012. Bet-hedging -a triple trade-off between means, variances and correlations. Biol. Rev. 87:742–755.

Stephens, D. W. 2007. Models of information use. Pp. 31–60 in D. W. Stephens, J. S. Brown, and R. C. Ydenberg, eds. Foraging. University of Chicago Press, Chicago.

Tarka, M., B. Hansson, and D. Hasselquist. 2015. Selection and evolutionary potential of spring arrival phenology in males and females of a migratory songbird. J. Evol. Biol. 28:1024–1038.

Thorley, J. B., and A. M. Lord. 2015. Laying date is a plastic and repeatable trait in a population of blue tits *Cyanistes caeruleus*. Ardea 103:69–78.

Tufto, J. 2015. Genetic evolution, plasticity, and bet-hedging as adaptive responses to temporally autocorrelated fluctuating selection: A quantitative genetic model. Evolution 69:2034–2049.

Van Buskirk, J. 2012. Behavioural plasticity and environmental change. Pp. 145–158 in U. Candolin and B. B. M. Wong, eds. Behavioural Responses to a Changing World. Oxford University Press, Oxford.

Van Buskirk, J., and U. K. Steiner. 2009. The fitness costs of developmental canalization and plasticity. J. Evol. Biol. 22:852–860.

Venable, D. L., and J. S. Brown. 1988. The selective interactions of dispersal, dormancy, and seed size as adaptations for reducing risk in variable environments. Am. Nat. 131:360–384.

Verhulst, S., and J. Å. Nilsson. 2008. The timing of birds’ breeding seasons: A review of experiments that manipulated timing of breeding. Philos. Trans. R. Soc. B Biol. Sci. 363:399–410.

Verhulst, S., J. H. van Balen, and J. M. Tinbergen. 1995. Seasonal decline in reproductive success of the Great tit: Variation in time or quality? Ecology 76:2392–2403.

Via, S., R. Gomulkiewicz, G. De Jong, S. M. Scheiner, C. D. Schlichting, and P. H. Van Tienderen. 1995. Adaptive phenotypic plasticity: consensus and controversy. Trends Ecol. Evol. 10:212–217.

Visser, M. E., C. Both, and M. M. Lambrechts. 2004. Global climate change leads to mistimed avian reproduction. Adv. Ecol. Res. 35:89–110.

Wang, C.-C., and D. C. Rogers. 2018. Bet hedging in stochastic habitats: an approach through large branchiopods in a temporary wetland. Oecologia 188:1081–1093. Springer Berlin Heidelberg.

Westneat, D. F., J. Wright, and N. J. Dingemanse. 2015. The biology hidden inside residual within-individual phenotypic variation. Biol. Rev. 90:729–743.

Wright, J., G. H. Bolstad, Y. G. Araya-Ajoy, and N. J. Dingemanse. 2019. Life-history evolution under fluctuating density-dependent selection and the adaptive alignment of pace-of-life syndromes. Biol. Rev. 94:230–247.

Xue, B., P. Sartori, and S. Leibler. 2019. Environment-to-phenotype mapping and adaptation strategies in varying environments. Proc. Natl. Acad. Sci. 116:13847–13855.

Ydenberg, R. C., J. S. Brown, and D. W. Stephens. 2007. Foraging: An overview. Pp. 1–28 in D. W. Stephens, J. S. Brown, and R. C. Ydenberg, eds. Foraging: Behavior and ecology. University of Chicago Press, London.

Yoshimura, J., and C. W. Clark. 1991. Individual adaptations in stochastic environments. Evol. Ecol. 5:173–192.

